# Prefrontal stimulation disrupts motor memory consolidation at the micro timescale

**DOI:** 10.1101/2022.11.01.514668

**Authors:** Mareike A. Gann, Nina Dolfen, Bradley R. King, Edwin M. Robertson, Geneviève Albouy

## Abstract

Functional brain responses in hippocampo- and striato-cortical networks during initial motor sequence learning (MSL) are critical for memory consolidation. We have recently shown that prefrontal stimulation applied prior to initial MSL can alter these learning-related responses. In the present study, we investigated whether such stimulation-induced modulations of brain responses can influence motor memory consolidation at different timescales. Specifically, we examined the effect of prefrontal stimulation on the behavioral and neural responses associated to (i) fast consolidation processes occurring during short rest episodes interspersed with practice during initial learning (i.e., micro timescale) and (ii) slow consolidation process taking place across practice sessions separated by 24h (i.e., macro timescale). To do so, we applied active (inhibitory or facilitatory) or control theta-burst stimulation to the prefrontal cortex of young healthy participants before they were trained on an MSL task while their brain activity was recorded using functional magnetic resonance imaging (fMRI). Motor performance was retested, in the MRI scanner, after a night of sleep. Both our behavioral and brain imaging results indicate that while stimulation did not modulate consolidation at the macro timescale, it disrupted the micro-offline consolidation process. Specifically, our behavioral data indicate that active - as compared to control - stimulation resulted in a decrease in micro-offline gains in performance over the short rest intervals. At the brain level, stimulation disrupted activity in the caudate nucleus and the hippocampus during the micro-offline intervals. Additionally, multivariate pattern persistence from task into inter-practice rest episodes - which is thought to reflect the reactivation of learning-related patterns - was hindered by active prefrontal stimulation in the hippocampus and caudate nucleus. Importantly, stimulation also altered the link between the brain and the behavioral markers of the micro-offline consolidation process. These results collectively suggest that active prefrontal stimulation prior to MSL disrupted both the behavioral and neural correlates of motor memory consolidation at the micro timescale.

## 1 Introduction

Motor memory consolidation is the offline (i.e., in the absence of task practice) process by which novel motor memory traces are reorganized into stabile representations (Robertson et al., 2004). Traditionally, consolidation processes have been assessed at the macro timescale (i.e., hours to days following initial learning). This research has demonstrated that macro-offline consolidation is facilitated by sleep; specifically, motor memory traces are less susceptible to interference from competing material and performance levels are maintained or even improved after a sleep period (e.g., (Robertson et al., 2004; Albouy et al., 2013a, 2013b, 2015); for an extensive review see (King et al., 2017)). At the brain level, responses in the hippocampus as well as competitive interactions between striato-cortical and hippocampo-cortical circuits during initial motor sequence learning (MSL) have been shown to predict the subsequent sleep-related macro-offline consolidation process (Albouy et al., 2008, 2013a, 2013b). Interestingly, recent research has shown that consolidation can also occur on a faster timescale (i.e., seconds to minutes) during the short rest periods interspersed with practice (Bönstrup et al., 2019). During these micro-offline intervals, gains in performance are observed and the corresponding fast consolidation process is also associated with brain responses in the hippocampus (Jacobacci et al., 2020; Buch et al., 2021). The evidence described above highlights the central role of the hippocampus in motor memory consolidation at both micro and macro timescales. Importantly, it remains uncertain whether altering these neural signatures with experimental interventions can effectively influence consolidation. In line with a series of hippocampal-targeted non-invasive brain stimulation studies in the declarative memory domain (e.g., (Wang et al., 2014; Kim et al., 2018; Tambini et al., 2018; Freedberg et al., 2019; Hermiller et al., 2019, 2020; Warren et al., 2019)), we recently showed that learning-related responses in the hippocampus were influenced by stimulation of the dorsolateral prefrontal cortex (DLPFC) applied prior to initial MSL (Gann et al., 2021b, 2021a). Specifically, inhibitory - as compared to facilitatory - thetaburst stimulation (TBS) of the DLPFC altered functional connectivity patterns in fronto-hippocampal networks over the course of learning. The intervention also modulated task-related striatal responses which are known to play a critical role in the motor learning and memory consolidation process (Albouy et al., 2013a). Importantly, no studies have ever investigated whether such stimulation-induced modulations of brain responses during initial learning can influence the behavioral and neural markers of consolidation at the micro and/or macro timescales.

To address this question, we applied inhibitory continuous (cTBS), facilitatory intermittent (iTBS) or control theta-burst stimulation to the DLPFC of young healthy participants before they were trained on an MSL task while their brain activity was recorded using functional magnetic resonance imaging (fMRI). Motor performance was retested in the MR scanner after a night of sleep (see Figure 1). We combined univariate and multivariate analyses of task- and rest-related fMRI data to comprehensively characterize the effect of prefrontal stimulation on the neural processes supporting motor memory consolidation at different timescales. Based on our earlier work (Gann et al., 2021a), we expected inhibitory cTBS to specifically alter fronto-hippocampal as well as striatal responses during initial MSL as compared to iTBS and control stimulation. As these brain responses during initial training are linked to consolidation process at both the micro (Jacobacci et al., 2020; Buch et al., 2021) and macro (Albouy et al., 2008, 2013a, 2013b) timescales, we expected cTBS to compromise consolidation at both timescales. Brain responses during retest were expected to be indirectly influenced by the effect of stimulation during initial learning. Specifically, as a consequence of altered striatal and hippocampal responses during initial training under cTBS, we predicted that the motor memory trace during retest would remain supported by brain areas involved in early learning stages (e.g., hippocampus and associative regions of the striatum).

**Figure 1.**
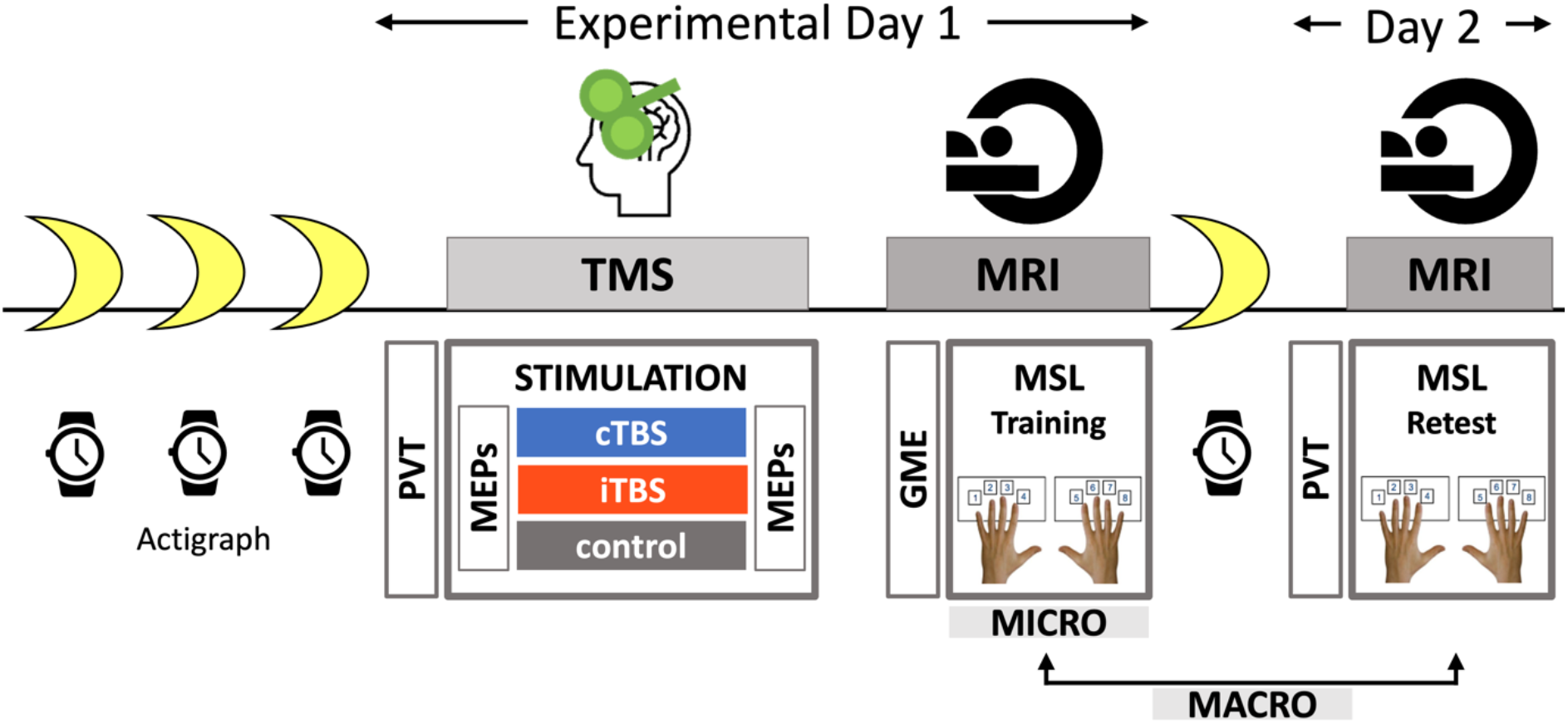
Experimental design. Participants were instructed to follow a regular sleep schedule 3 nights before experimental day 1 until the end of the experiment (4 nights total, compliance checked with actigraphy and sleep diary). On experimental day 1, theta-burst stimulation (TBS) was applied to the dorsolateral prefrontal cortex (DLPFC, −30 22 48mm) prior to motor sequence learning (MSL). Corticospinal excitability of the primary motor cortex was measured with motor evoked potentials (MEPs) pre- and post-TBS. Immediately following stimulation, participants were placed in the magnetic resonance imaging (MRI) scanner where general motor execution (GME) was probed with a random serial reaction time task immediately prior training on the MSL task. On the next day (day 2), participants were retested on the MSL task in the MRI scanner, approximately 24h after initial training. The night of sleep between experimental days 1 and 2 was spent at home and monitored with sleep diary and actigraphy. Objective vigilance was assessed at the start of each session with a psychomotor vigilance test (PVT) on both experimental days. This design allowed us to examine the effect of prefrontal stimulation on the behavioral and neural responses associated to (i) fast consolidation processes occurring during short rest episodes interspersed with practice during initial learning (i.e., MICRO timescale) and (ii) slow consolidation process taking place across practice sessions separated by 24h (i.e., MACRO timescale). TMS: transcranial magnetic stimulation, cTBS: continuous theta-burst stimulation, iTBS: intermittent theta-burst stimulation.

## 2 Results

This study was pre-registered in the Open Science Framework (https://osf.io/e2cnq). Analyses reported below that were not included in the pre-registration are labeled as exploratory.

### 2.1 Behavioral analyses

#### 2.1.1. Initial learning session

After receiving active or control stimulation to the prefrontal cortex, participants learned a bimanual sequence of eight finger movements through repeated practice (Figure 1; MSL training; 20 practice blocks alternating with 15s rest blocks). Behavioral results show that both performance speed (inter-trial intervals) and accuracy (i.e., percentage of correct transitions) significantly improved across practice blocks during the MSL training session (see Figure 2A, training panel: blocks 1-20; main effect of block: speed, F_(6.798,448.694)_=73.471, η_p_^2^=0.527, *p*<0.001; accuracy, F_(12.331,813.87)_=1.811, η_p_^2^=0.027, *p*=0.041) similarly between groups (block by group interaction: speed, F_(13.597,448.694)_=0.933, η_p_^2^=0.028, *p*=0.587; accuracy, F_(24.663,813.87)_=0.854, η_p_^2^=0.025, *p*=0.721). While performance was overall faster during training in the cTBS group as compared to the other groups, the group effect was not significant (speed, F_(2,66)_=0.65, η_p_^2^=0.024, *p*=0.444; accuracy, F_(2,66)_=0.13, η_p_^2^=0.004, *p*=0.878). These results suggest that prefrontal stimulation prior to MSL did not influence overall motor performance during initial training.

**Figure 2.**
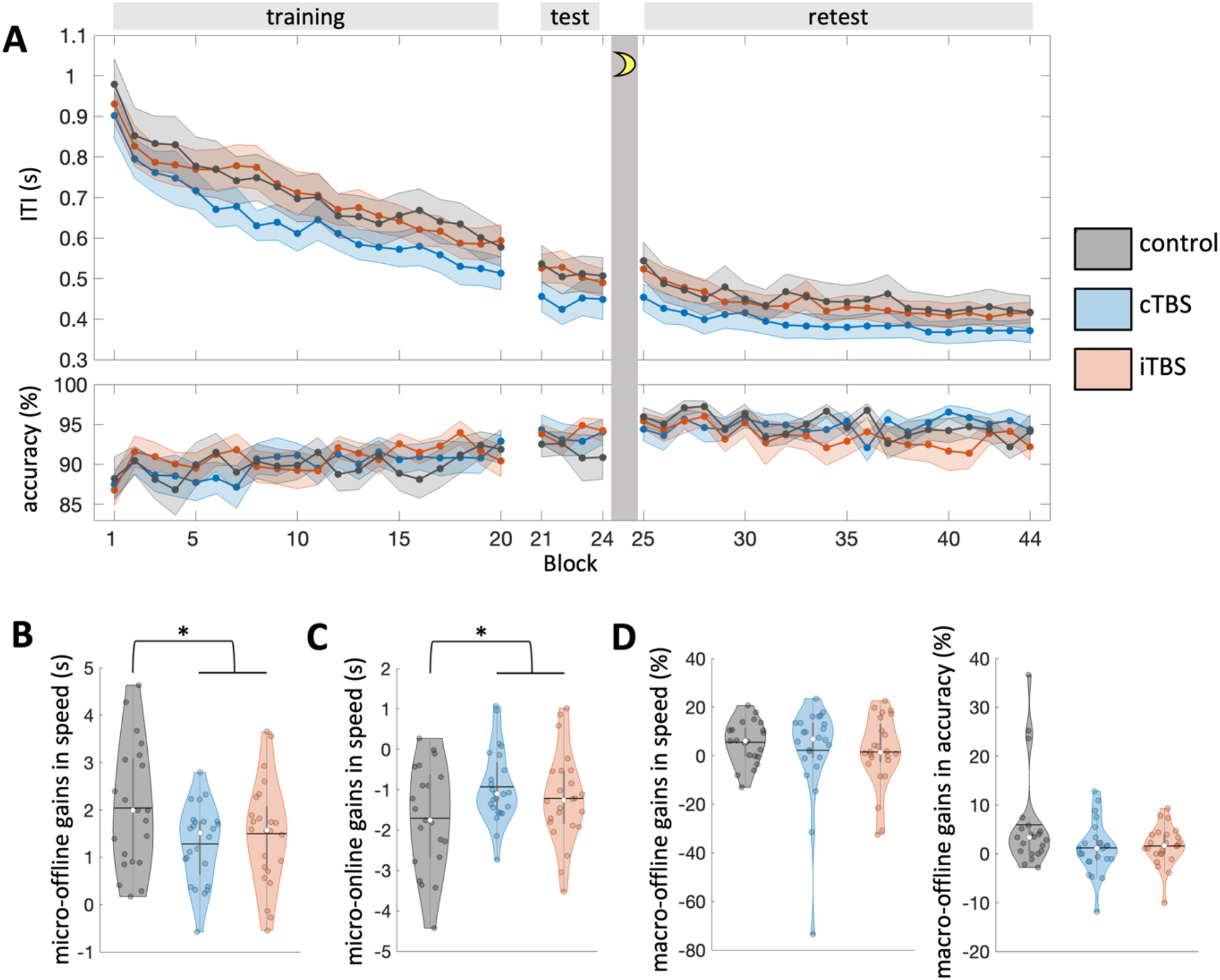
Performance speed and accuracy. (A) Both performance speed (intertrial interval, ITI, in s) and accuracy (correct transitions in %) improved over the course of initial training (first 20 blocks). Accuracy remained high and stable during the subsequent sessions. In contrast, further improvements were observed for performance speed during the test and retest sessions (block 21-24 and 25-44, respectively, see Supplemental Results). No main effects of group were observed in any of the practice sessions. Dots represent mean values; shaded areas represent standard errors of the mean. While overall learning (A) did not differ among the three groups, active - as compared to control - stimulation resulted in (B) lower micro-offline and (C) higher micro-online gains in performance speed (in s). (D) Overnight macro-offline gains in performance speed (%) did not differ between groups (left panel) but a significant group effect was observed on macro-offline gains in performance accuracy whereby gains were higher in control as compared to the TBS groups (right panel). Note that this effect was driven by extreme values in the control group. In panels B-D, colored circles represent individual data, jittered in arbitrary distances on the x-axis within the respective violin plot to increase perceptibility. Black horizontal lines represent means and white circles represent medians. The shape of the violin plots depicts the distribution of the data and grey vertical lines represent quartiles. Violin plots were created with (Bechtold, 2016). cTBS: continuous theta-burst stimulation, iTBS: intermittent theta-burst stimulation.

We then performed exploratory analyses to examine the effect of stimulation on the fast consolidation process occurring at the micro timescale with procedures similar to previous research (Bönstrup et al., 2019; Jacobacci et al., 2020; Buch et al., 2021). Specifically, we tested whether stimulation influenced (i) *micro-offline gains in performance speed* defined as the performance changes across the short rest intervals, i.e., from the end of one to the beginning of the next practice block, as well as (ii) *micro-online gains in performance speed* computed as the performance changes from the beginning to the end of a practice block during initial learning (see Methods for further information). Results showed a trend for a group main effect on micro-offline gains in performance (F_(2,66)_=2.904, η_p_^2^=0.081, *p*=0.062). Inspection of the data and follow-up comparisons (Supplemental Table S1) indicated a similar pattern of results in the two active stimulation groups as compared to control. Follow-up analyses collapsing across iTBS and cTBS groups revealed that active - as compared to control - stimulation resulted in decreased micro-offline gains in performance (active vs. control; *t*_(29.685)_=-2.05, d=-0.606, *p*=0.049; Figure 2B). Micro-online gains in performance also showed a trend for a group effect (F_(2,66)_=2.786, η_p_^2^=0.078, *p*=0.069) with greater micro-online gains in the active as compared to the control stimulation group (active vs. control; *t*_(67)_=2.19, d=0.573, *p*=0.032; Figure 2C; see Supplemental Table S1 for group pair comparisons). Altogether, these results suggest that active DLPFC stimulation applied prior to MSL altered the balance between micro-online and -offline processes during initial learning such that online and offline learning were respectively enhanced and disrupted as compared to control stimulation.

#### 2.1.2. Between-session changes in performance

We examined the effect of stimulation on the slow consolidation process occurring at the macro timescale, i.e., between practice sessions. To do so, overnight macro-offline gains in performance were computed as the percentage of the difference in performance from the MSL test (average of the 4 blocks) to the beginning of the MSL retest (average first 4 blocks) for performance speed and accuracy (see Supplemental Results for separate statistical analyses on MSL test and retest sessions). While no group effect was observed on macro-offline gains in performance speed (F_(2,66)_=0.426, η_p_^2^=0.013, *p*=0.655; Figure 2D, left panel), macro-offline gains in performance accuracy were significantly different among groups (F_(2,66)_=3.287, η_p_^2^=0.091, *p*=0.044; Figure 2D, right panel). However, post-hoc two-tailed two-sample *t* tests did *not* show significant differences between any pair of groups after Bonferroni correction (cTBS vs. iTBS: *t*_(46)_=-0.284, d=0.082, *p*=0.778, *p_Bonferroni_*=1; cTBS vs. control: *t*_(43)_=-2.012, d=-0.601, *p*=0.051, *p_Bonferroni_*=0.153; iTBS vs. control: *t*_(25.617)_=-1.859, d=-0.584, *p*=0.074, *p_Bonferroni_*=0.222). Also note that the effect seemed to be driven by a few extreme participants in the control group showing high accuracy gains. Altogether, these results indicate that prefrontal stimulation did not effectively modulate motor memory consolidation at the macro timescale.

In conclusion, our behavioral results indicate that prefrontal stimulation applied prior to motor sequence learning disrupted the micro-offline consolidation process but did not affect consolidation at the macro timescale.

### 2.2 Functional brain imaging data analyses

The fMRI analyses presented below employed a region of interest (ROI) approach to examine the effect of prefrontal stimulation on brain responses in the TMS target (DLPFC), the hippocampus and the basal ganglia.

#### 2.2.1. Initial learning session

##### Univariate analyses

In a first set of univariate analyses of the fMRI data related to the initial MSL training session, we examined the effect of prefrontal stimulation applied prior to MSL on task-related brain activity (task vs. rest). Results showed a main effect of group in the hippocampus and the caudate nucleus (Table 1.1). Follow-up two-sample *t* tests (Supplemental Table S2) and data inspection (Figure 3A) indicated that this effect was driven by more deactivation (i.e., more activity during inter-practice rest intervals as compared to task practice) in the control group as compared to the two active stimulation groups (iTBS and cTBS groups). Exploratory analyses pooling the two active stimulation groups together confirmed this pattern of results (Table 1.2). Altogether, these data suggest that active, as compared to control, stimulation disrupted hippocampal and caudate activity during inter-practice rest (as compared to task) intervals of the initial training session.

**Figure 3.**
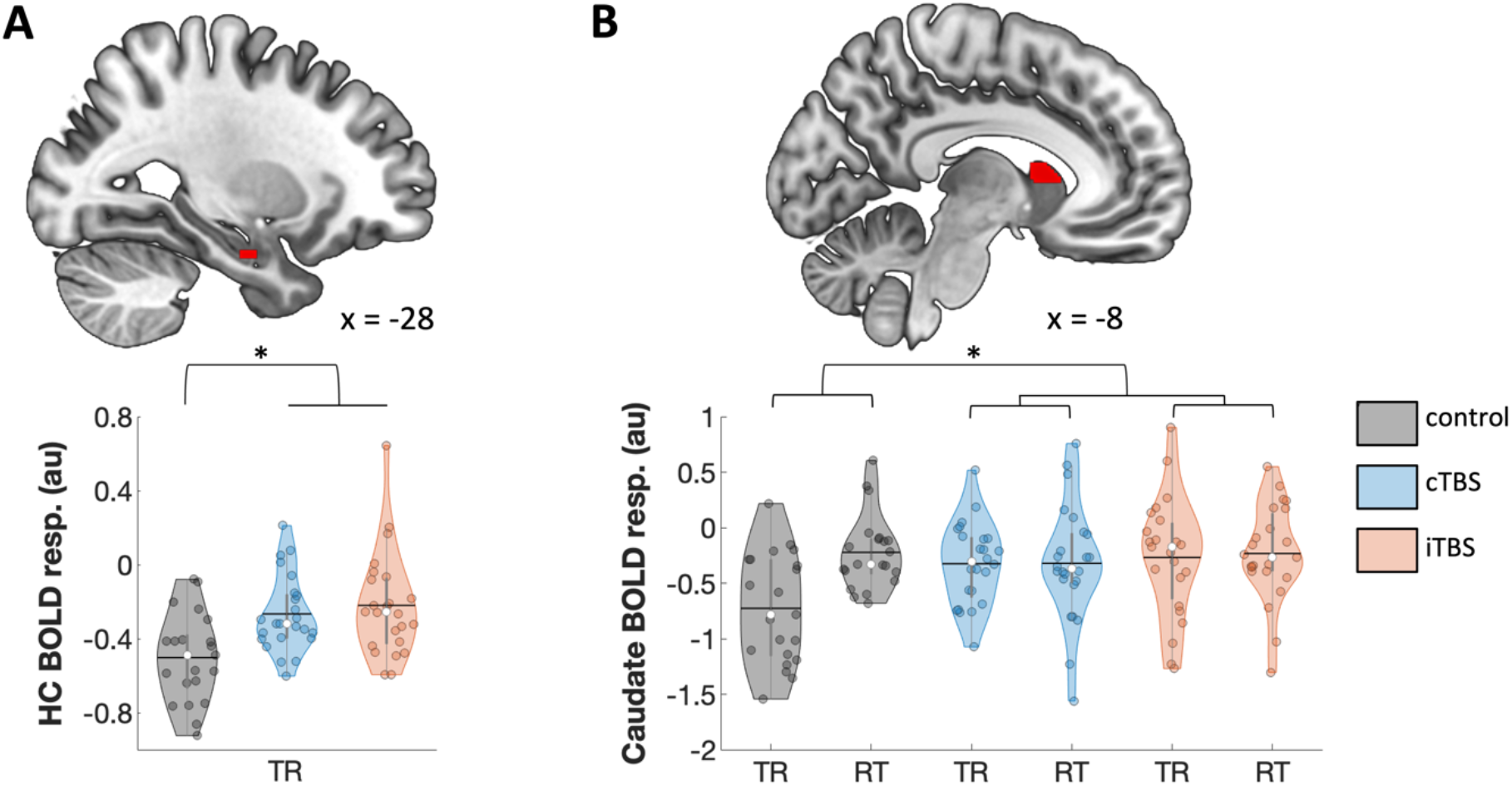
Main effect of group on brain activity. (A) MSL training (TR). The group effect observed in the hippocampus (HC) was explained by greater task-related deactivation (i.e., higher activity during inter-practice rest intervals) in the control group compared to the two active TBS groups. (B) MSL training (TR) – retest (RT). The group effect observed in the caudate was driven by larger between-session changes in activity in the control group (more deactivation during training than during retest) as compared to the two active TBS groups. Activation maps are displayed within the ROIs on a T1-weighted template image at a threshold of *p*<0.005, uncorrected. Asterisks indicate significant group differences between group pairs *(p_FWE_svc__*<0.05, see Supplemental Table S1 for paired comparisons). Colored circles represent individual data, jittered in arbitrary distances on the x-axis within the respective violin plot to increase perceptibility. Black horizontal lines represent means and white circles represent medians. The shape of the violin plots depicts the distribution of the data and grey vertical lines represent quartiles. Resp.: response, au: arbitrary unit, cTBS: continuous theta-burst stimulation, iTBS: intermittent theta-burst stimulation.

**Table 1.** Functional imaging results of activation-based contrasts (Main effect of practice)

| Area | x | y | z | k | Z | $p_{FWESvc}$ |
| --- | --- | --- | --- | --- | --- | --- |
| **1. Main effect of group (F test)** |  |  |  |  |  |  |
| **TRAINING** |  |  |  |  |  |  |
| Hippocampus | -28 | -10 | -26 | 16 | 3.1 | 0.027 |
| Caudate | -8 | 16 | 4 | 47 | 2.97 | 0.037 |
| **RETEST** |  |  |  |  |  |  |
| No significant responses |  |  |  |  |  |  |
| **TRAINING - RETEST** |  |  |  |  |  |  |
| Caudate | -6 | 12 | 6 | 26 | 2.94 | 0.042* |
| **2. Main effect of active stimulation ([iTBS+cTBS] vs. control) (t test)** |  |  |  |  |  |  |
| **TRAINING** |  |  |  |  |  |  |
| **[control-active]** |  |  |  |  |  |  |
| No significant responses |  |  |  |  |  |  |
| **[active-control]** |  |  |  |  |  |  |
| Hippocampus | -28 | -10 | -26 | 55 | 3.67 | 0.0038* |
| Caudate | -8 | 16 | 4 | 47 | 3.58 | 0.0051* |
| Putamen/Pallidum | -10 | 0 | 6 | 175 | 2.88 | 0.036 |
| Pallidum | -12 | -2 | -2 | 23 | 3.09 | 0.021 |
| Caudate | 8 | 14 | 4 | 31 | 2.83 | 0.041 |
| **RETEST** |  |  |  |  |  |  |
| **[control-active]** |  |  |  |  |  |  |
| No significant responses |  |  |  |  |  |  |
| **[active-control]** |  |  |  |  |  |  |
| Caudate | -12 | 14 | -6 | 19 | 3.02 | 0.024* |
| **TRAINING - RETEST** |  |  |  |  |  |  |
| **[control-active]** |  |  |  |  |  |  |
| No significant responses |  |  |  |  |  |  |
| **[active-control]** |  |  |  |  |  |  |
| Caudate | -8 | 10 | 10 | 87 | 3.71 | 0.003* |

Asterisk (\*) indicates significance at $p < .05$ after Holm-Bonferroni correction for multiple comparison.

##### Multivariate analyses

Both the behavioral and univariate fMRI analyses reported above suggest that stimulation altered processes occurring during the micro-offline rest epochs. Interestingly, recent MEG research suggests that task-relevant brain activity is reactivated (replayed) during these inter-practice rest intervals (Buch et al., 2021). An approach that has shown promise to study such replay using fMRI data is multivariate pattern analyses. Specifically, multivoxel correlation structure (MVCS) analyses allow the examination of the persistence of task patterns into subsequent rest; a process that is thought to reflect the reactivation of learning-related patterns during offline epochs (Tambini and Davachi, 2013; Gann et al., 2021b; King et al., 2021). Thus, in a second set of exploratory fMRI analyses, we used MVCS approaches to test whether stimulation influenced the persistence of task patterns into inter-practice rest intervals. To do so, we computed the level of similarity (similarity index) of multi-voxel correlation structures between task practice and inter-practice rest periods (see Figure 4A) in the two ROIs highlighted above (caudate nucleus and hippocampus) and in a control subcortical region (thalamus). Results of these analyses showed that the persistence of task patterns into inter-practice rest epochs did not differ among the three groups in the caudate nucleus and the hippocampus (caudate nucleus: F_(2,66)_=1.811, η_p_^2^=0.052, *p*=0.171; hippocampus: F_(2,66)_=1.1, η_p_^2^=0.032, *p*=0.339). As above, analyses collapsing the two active stimulation groups together indicated that similarity indices between task and rest were higher in the control as compared to the active stimulation conditions in the caudate nucleus (*t*_(67)_=1.899, d=-0.497, *p*=0.031; see Supplemental Table S3 for all pair-wise comparisons; Figure 4B). A similar trend was observed in the hippocampus (*t*_(67)_=1.494, d=-0.391, *p*=0.07; Figure 4C). Interestingly, no such effects were observed in the control subcortical region (thalamus; group effect: F_(2,66)_=0.82, η_p_^2^=0.024, *p*=0.445; active vs. control stimulation: *t*_(67)_=1.088, d=0.285, *p*=0.14; Figure 4D). In sum, the MVCS results indicate that active stimulation disrupted the persistence of task-related patterns into the inter-practice rest periods in the caudate nucleus and, to a lesser extent, the hippocampus. These findings suggest that DLPFC stimulation applied prior to MSL hindered the reactivation of learning-related patterns during the micro-offline rest episodes.

**Figure 4.**
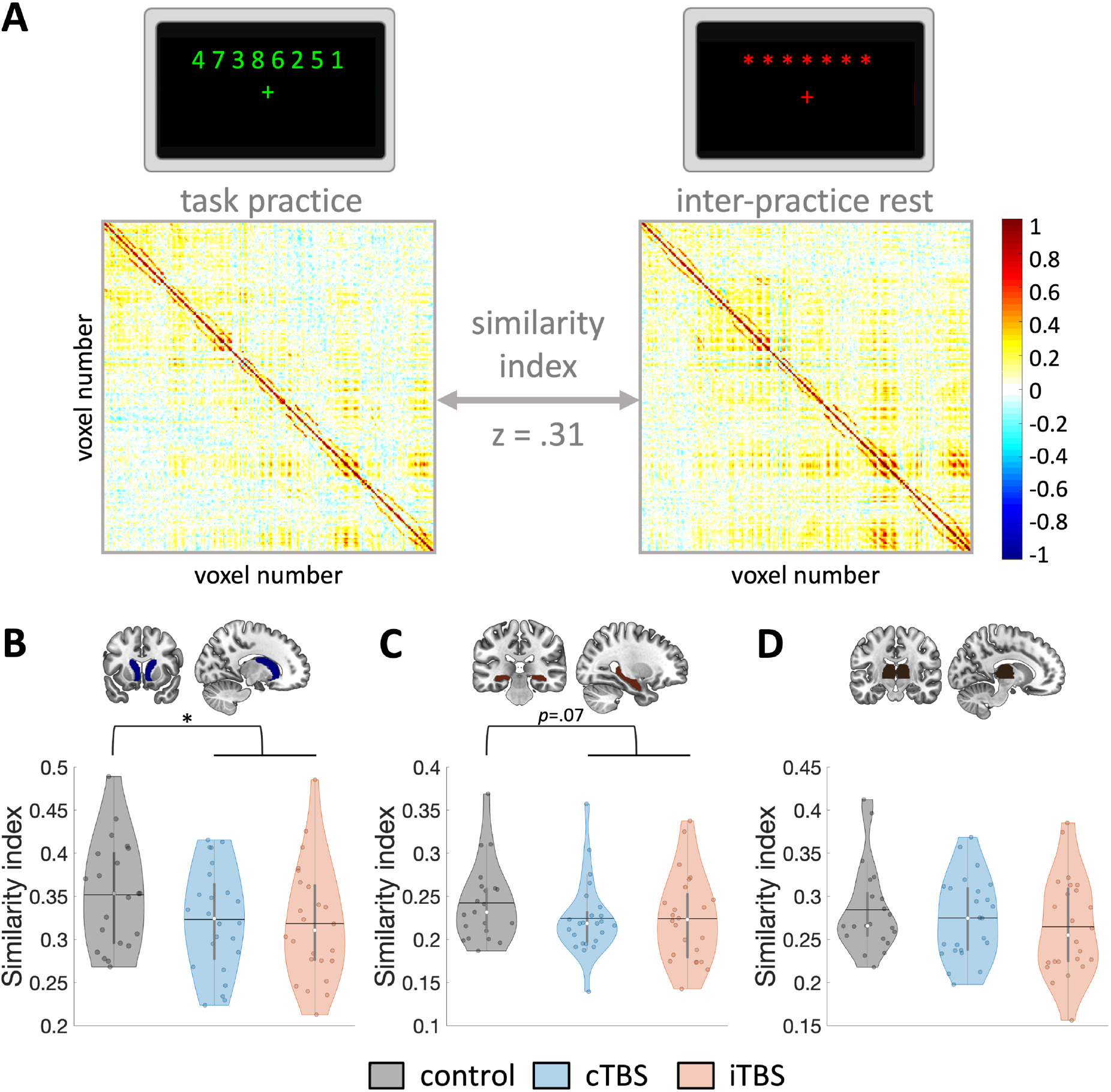
Similarity index between task practice and inter-practice rest periods during initial MSL. (A) Multivoxel correlation structure (MVCS) for an exemplar ROI and participant. Each matrix depicts the correlation between each of the n voxels of the ROI with all the other voxels of the ROI during task and inter-practice rest periods. The similarity between two matrices is calculated as the r-to-z transformed correlation between the two MVCS. Resulting Z scores are compared between stimulation groups. (B) The similarity index, reflecting the persistence of brain patterns from task practice into inter-practice rest periods, was lower in the active (cTBS and iTBS) groups as compared to the control stimulation in the caudate nucleus. (C) A similar effect was trending for the hippocampus (*p*=0.07), but not for (D) the thalamus. Colored circles represent individual data, jittered in arbitrary distances on the x-axis within the respective violin plot to increase perceptibility. Black horizontal lines represent means and white circles represent medians. The shape of the violin plots depicts the distribution of the data and grey vertical lines represent quartiles. Violin plots were created with (Bechtold, 2016). ROI masks are depicted on a T1-weighted template image. cTBS: continuous theta-burst stimulation, iTBS: intermittent theta-burst stimulation.

##### BOLD amplitude / pattern persistence relationships

The fMRI analyses reported above suggest that active prefrontal stimulation modulated both the amplitude of the BOLD response and the persistence of task patterns during inter-practice rest epochs. To test whether these metrics were related, we performed exploratory correlation analyses between the amplitude of the BOLD response in the ROIs (i.e., beta estimates extracted from the univariate fMRI analyses reported above, see Table 1.1) and pattern persistence (similarity index) measured with the multivariate approach; and we examined whether stimulation conditions altered this relationship. Results show that there was no main effect of group (χ^2^=0.33, *p*=0.85) or difference between active and control stimulation conditions (χ^2^=0.23, *p*=0.629) in the relationship between activity and pattern persistence in the caudate nucleus (see Supplemental Table S4 for results in each group). However, in the hippocampus, this relationship differed among the three groups (χ^2^=7.68, *p*=0.021, see Supplemental Table S4 for group pair comparisons) as well as between the active and control stimulation conditions (χ^2^=5.98, *p*=0.014; Figure 5A). Specifically, higher hippocampal activity during inter-practice rest blocks (i.e., stronger deactivation during practice blocks) was related to higher pattern persistence in the control group as compared to the active stimulation groups. These results not only suggest that the amplitude of the BOLD signal in the hippocampus during inter-practice rest periods is related to pattern persistence (and might therefore reflect reactivation of learning-related patterns), but also that active DLPFC stimulation disrupted this relationship.

**Figure 5.**
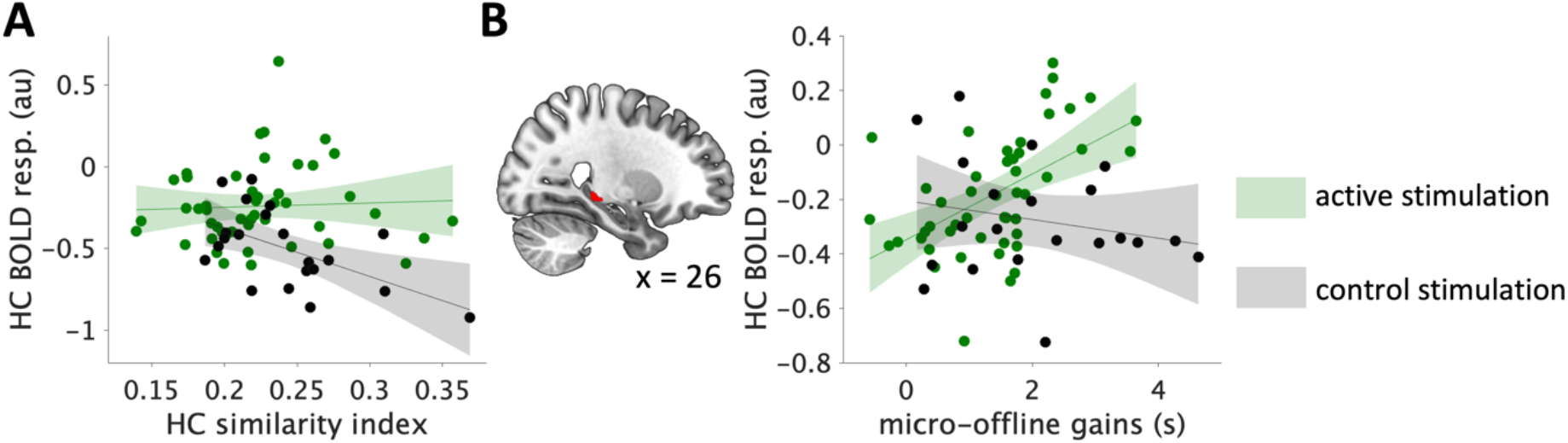
(A) Active (cTBS and iTBS) vs. control stimulation group differences in the relationship between task-related BOLD responses (beta values extracted from the hippocampal coordinate reported in Table 1.1) and similarity of hippocampal patterns between task and inter-practice rest during initial MSL. Higher hippocampal activity during inter-practice rest intervals (i.e., more negative beta values) was related to higher persistence (reflected by higher similarity index) of brain patterns from task practice into rest epochs in the control group compared to the active stimulation groups. (B) Regression analysis showing active vs. control stimulation group difference in the relationship between task-related responses in the hippocampus and micro-offline gains in performance speed during initial MSL. Higher hippocampal activity during inter-practice rest intervals (i.e., more negative beta values) was related to greater micro-offline gains in performance in the control group and lower gains in the active stimulation groups. Regression maps are displayed in the ROIs on a T1-weighted template image at a threshold of *p*<0.005, uncorrected. Circles represent individual data, solid lines represent linear regression fits, shaded areas depict 95% prediction intervals of the linear function. HC: Hippocampus, resp.: response, au: arbitrary unit.

##### Brain / behavior relationships

We conducted exploratory analyses to test whether there was a relationship between the behavioral and brain markers of the micro-offline consolidation processes and whether this relationship was influenced by stimulation. First, we examined BOLD / behavior relationships using micro-offline gains in performance as covariate in an univariate regression analyses. Similar as above, while the relationship between BOLD responses and micro-offline gains in performance did not differ among the three groups in any of the ROIs, this relationship differed between active and control stimulation conditions in the hippocampus (Table 2 and Supplemental Table S5; Figure 5B). Specifically, higher hippocampal activity during inter-practice rest blocks was related to higher micro-offline gains in performance in the control group and lower gains in the active stimulation groups. These findings suggest that active stimulation modulated the relationship between brain and behavioral markers of the micro-offline memory consolidation process. Second, we tested whether pattern persistence metrics were related to the amplitude of the micro-offline gains in performance. Results did not reveal any significant group effect or active vs. control stimulation effect in the ROIs (group comparison: caudate nucleus: χ^2^=2.49, *p*=0.288; hippocampus: χ^2^=0.18,*p*=0.913; thalamus: χ^2^=2.21, *p*=0.33; active vs. control stimulation: caudate nucleus: χ^2^=0.004, *p*=0.948; hippocampus: χ^2^=0.02, *p*=0.88; thalamus: χ^2^=0.3, *p*=0.583; see Supplemental Table S4 for group pair comparisons).

**Table 2.** Functional imaging results of the regression analyses between task-related activity maps (main effect of practice) and the sum of micro-offline gains during initial MSL

| Area | x | y | z | k | Z | $p_{FWEsvc}$ |
| --- | --- | --- | --- | --- | --- | --- |
| **1. Main effect of group (F test)** |  |  |  |  |  |  |
| No significant responses |  |  |  |  |  |  |
| **2. Active vs control stimulation (t tests)** |  |  |  |  |  |  |
| **[control-active]** |  |  |  |  |  |  |
| No significant responses |  |  |  |  |  |  |
| **[active-control]** |  |  |  |  |  |  |
| Hippocampus | 26 | -30 | -6 | 16 | 3.07 | 0.023* |

Altogether, the results of the fMRI data analyses related to the initial learning session indicate that active prefrontal - as compared to control - stimulation disrupted (1) brain responses reflecting motor memory reactivation during the micro-offline episodes and (2) the link between the brain and behavioral markers of the micro-offline motor memory consolidation process.

#### 2.2.2. Between-session changes in brain activity

We examined the effect of prefrontal stimulation on the changes in brain activity occurring between the training and the overnight retest sessions which indirectly reflect the slower consolidation process taking place at the macro timescale. Results indicate a main effect of group on the changes in brain activity (task vs. rest) from training to retest in the caudate nucleus (Table 1.1). Specifically, between-session changes in caudate activity were larger in the control group than in the two active stimulation groups. These effects were explained by the patterns of deactivation observed in the control group during training. Specifically, there was more deactivation during inter-practice rest intervals during training as compared to retest in the control group as compared to the two active stimulation groups (Figure 3B, Table 1.2, Supplemental Table S2). In line with these results, there was no main effect of group on brain activity (task vs. rest) during the overnight retest session.

Altogether, these results suggest that prefrontal stimulation did not influence the changes in brain activity occurring as a result of the macro-offline memory consolidation process.

## 3 Discussion

In the present study, we investigated whether prefrontal stimulation prior to initial motor sequence learning altered consolidation at different timescales (i.e., micro and macro). Our results indicated that while stimulation did not modulate the macro-offline consolidation process, it disrupted micro-offline consolidation occurring on a faster timescale. Specifically, active stimulation resulted in a decrease in micro-offline gains in performance observed over the short inter-practice rest intervals during initial learning. At the brain level, stimulation disrupted activity in the caudate nucleus and the hippocampus during these micro-offline intervals. Additionally, multivariate pattern persistence from task into inter-practice rest episodes - which is thought to reflect the reactivation of learning-related patterns - was hindered by active prefrontal stimulation in the same brain regions (i.e., hippocampus and caudate nucleus). Importantly, our results also show that stimulation altered the link between the behavioral and brain markers of the micro-offline consolidation process.

### Prefrontal stimulation altered consolidation processes at the micro timescale

Earlier research has consistently shown that motor memory consolidation also occurs at a fast timescale during the short micro-offline rest intervals interspersed with task practice (Bönstrup et al., 2019, 2020; Jacobacci et al., 2020; Buch et al., 2021; Quentin et al., 2021). These studies have collectively demonstrated that overall gains in performance during initial motor sequence learning are primarily driven by micro-offline gains occurring across inter-practice rest intervals, as limited improvement in performance is observed during online task practice. Our behavioral data confirm these previous findings. Importantly, our analyses also showed that active stimulation modulated these micro timescale consolidation processes. Specifically, DLPFC stimulation applied prior to MSL altered the balance between micro-online and micro-offline gains in performance such that online and offline learning were respectively enhanced and disrupted as compared to control stimulation. Together with the results of both the univariate and multivariate analyses of our neuroimaging data suggesting a disruption of the brain responses associated to micro-offline processes, we speculate that greater micro-online gains in performance under stimulation might compensate for the disruption of the micro-offline processes. These compensatory processes might ultimately result in similar overall performance between the active stimulation and control groups.

Recent research has associated hippocampal responses during inter-practice rest intervals to micro-offline consolidation (Jacobacci et al., 2020; Buch et al., 2021). Our brain imaging data analyzed with both univariate and multivariate approaches are collectively in line with these earlier observations but also suggest that active DLPFC stimulation disrupted hippocampal responses during inter-practice rest. Interestingly, similar results were observed in the caudate nucleus, a brain region which, together with the hippocampus, is part of an associative network recruited early during training and supporting the abstract representation of the motor sequence (Albouy et al., 2013a). We speculate that the prefrontal stimulation-induced changes in brain responses in deep brain regions like the hippocampus and the caudate nucleus occurred via a modulation of connectivity in associative functional networks during learning (see (Gann et al., 2021a) and Supplemental Results 2.1.8 showing stimulation-induced modulation of DLPFC-caudate connectivity). Additionally, our data indicated that stimulation altered the relationship between hippocampal activity during these inter-practice rest periods (as compared to task) and micro-offline gains in performance such that greater hippocampal activity during rest was related to higher micro-offline gains in the control group and lower gains in the active stimulation groups. The brain-behavior relationship observed in the control group is in line with previous findings showing a positive correlation between hippocampal BOLD responses during rest and the amplitude of micro-offline gains in performance during early learning (Jacobacci et al., 2020). Interestingly, our data show that stimulation modulated the relationship between rest-related hippocampal activity and micro-offline gains in performance.

Recent advances in multivariate analyses of fMRI data have allowed a better characterization of the physiological processes supporting offline memory consolidation (Tambini and Davachi, 2013; King et al., 2021). Specifically, the persistence of task-related fMRI patterns into subsequent rest periods is thought to reflect the reactivation or replay of the memory trace during the consolidation episode. Such reactivations have been observed in the striatum and the hippocampus at the macro timescale, i.e., a few minutes after the end of initial motor learning (King et al., 2021). In the present study, we used this approach to examine the effect of stimulation on neuronal reactivation during the micro-offline consolidation period. The corresponding multivariate analyses showed that active prefrontal stimulation disrupted the persistence of task patterns into inter-practice rest periods in the caudate nucleus and the hippocampus. The current results suggest that hippocampal and caudate replays occur during the micro-offline consolidation episode - the former of which is in line with recent MEG findings (Buch et al., 2021) - and that this process is supported by similar neural mechanisms as during macro-offline episodes (i.e., reactivation of task patterns; (King et al., 2021)). Importantly, our data also show that prefrontal stimulation hindered the reactivation process in these deep brain regions that are critical for learning. Last, our findings provide evidence for a link between brain activity during rest and pattern persistence but also indicate that this relationship was disrupted by the stimulation. Specifically, higher hippocampal pattern persistence was related to greater hippocampal BOLD signal during inter-practice rest in controls more than in the stimulated groups. Altogether, the results of the multivariate analyses suggest that (1) task-related hippocampal and caudate patterns are reactivated during the micro-offline episodes, (2) the amplitude of the BOLD signal in the hippocampus during inter-practice rest periods reflects hippocampal reactivation, and (3) active prefrontal stimulation disrupted these processes.

### Prefrontal stimulation did not modulate consolidation at the macro timescale

Previous TMS studies have shown that DLPFC stimulation applied during or after motor learning can influence performance at a delayed retest (Galea et al., 2010; Ambrus et al., 2020). Based on these earlier studies and on our previous neuroimaging work showing that DLPFC stimulation can alter learning-related responses in hippocampal and striatal networks (Gann et al., 2021a) that are critical for the overnight consolidation processes (Albouy et al., 2008, 2013a, 2013b), we expected stimulation to modulate consolidation at the macro timescale. In contrast to our expectations, both our behavioral and brain imaging results indicate that DLPFC stimulation applied prior to MSL did not influence the subsequent sleep-related macro-offline consolidation process. It is important to mention though that the neural processes supporting macro-(in contrast to micro-) offline consolidation processes were only assessed indirectly in this study (i.e., contrasting brain activity between training vs. overnight retest sessions). The present data therefore do not allow us to conclude on the effect of stimulation on the neural processes supporting macro-offline consolidation *per se.* Nevertheless, we argue that the lack of stimulation effect on macro-offline consolidation might be related to the modulation of brain activity observed during learning. That is, the present intervention specifically altered hippocampal and striatal brain responses during inter-practice rest intervals, while brain activity described to predict overnight macro-offline gains are rather related to task practice (Albouy et al., 2008). Our results therefore suggest that stimulation-induced modulations of micro-offline consolidation did not affect the delayed macro-offline consolidation process. Interestingly, previous studies examining potential relationships between micro- and macro-offline memory processes have shown that micro-offline gains in performance are not related to gains observed overnight, i.e., at the macro timescale (Bönstrup et al., 2019; Jacobacci et al., 2020). In line with this research, our results indicated stimulation-induced modulations of consolidation at the micro but not the macro timescale which further substantiates the idea that these processes are independent (but see above for a discussion of the related brain responses). It is also worth noting that even though stimulation altered the balance between micro-online and micro-offline processes during initial learning, overall learning was not affected by the intervention. It is possible that macro-offline consolidation processes were therefore triggered irrespective of the nature of performance improvement during initial learning. Further research is warranted to better characterize possible interactions between micro- and macro-offline memory consolidation processes.

### Stimulation-induced effects did not differ between active stimulation groups

In contrast to our previous work showing a differential effect of iTBS vs. cTBS on task-related brain patterns and functional connectivity (Gann et al., 2021b, 2021a), the present analyses did not show such stimulation-specific effects. It is worth noting though that - similar to the present results - our previous studies did not highlight any effect of stimulation condition on brain *activity* during initial training (Gann et al., 2021a). One might argue that the effect of continuous and intermittent TBS on the DLPFC are less dichotomic (i.e., inhibition vs. facilitation) than those observed on M1 (Huang et al., 2005). This remains speculative as a direct measure of the effect of stimulation on DLPFC excitability - as done with MEP measurements after M1 stimulation - is not possible. However, evidence from earlier TMS-EEG studies does not support such view. Despite high inter-subject variability (Chung et al., 2019), plasticity processes in the DLPFC - as measured with electrophysiological responses - are described to be affected by TBS in a similar manner as in M1 (Chung et al., 2017, 2018b). It therefore remains unclear why the different stimulation conditions yielded similar results. However, the discrepancies with our previous work may be related to various factors such as the type of task used, the inclusion of a control task or not, as well as the study design. With respect to the latter point, our previous work made use of a within-subject design that typically provides higher statistical power to highlight more subtle effects as compared to the between-subject designs as used in our current study. Nevertheless, additional research is warranted to better characterize the specific effect of different frontal TBS protocols on brain functioning.

## 4 Conclusions

In the present study, we used hippocampal-network-targeted stimulation in order to modulate motor memory consolidation at different timescales. Altogether, our results indicated that while stimulation did not alter macro-offline consolidation, it hindered the faster micro-offline consolidation process. Specifically, we showed that prefrontal stimulation disrupted both the behavioral and brain markers of this fast plasticity processes as well as their relationship. This study provides the first evidence, to the best of our knowledge, that non-invasive-brain-stimulation of the prefrontal cortex can modulate fast motor memory consolidation through the modulation of deep brain responses during micro-offline episodes.

## 5 Methods

This study was pre-registered in the Open Science Framework (https://osf.io/e2cnq). Deviations from this protocol are marked in the main text with a number sign (#) and detailed in Supplemental Table S6 together with a justification for the change. Additionally, analyses that were not pre-registered are referred to as exploratory throughout the manuscript.

### 5.1 Ethics statement

Participants gave written informed consent before participating in this study. The current study was approved by the local Ethics Committee (UZ/KU Leuven) and was conducted according to the declaration of Helsinki (2013). All participants were compensated for their time and effort.

### 5.2 Participants

Seventy-six healthy young (age range 19-29 years, 52 females), right-handed (Edinburgh Handedness questionnaire, (Oldfield, 1971)) volunteers participated in the current study. One participant withdrew before group assignment and the 75 remaining participants were distributed in 3 experimental groups (control N=25; iTBS N=25; cTBS N=25). As per the preregistration, sample size estimation was based on detecting significant differences in the behavior of interest (i.e., macro-offline gains in performance, see below) among our three groups. A power analysis based on previous research (Debas et al., 2010) and using G*Power ((Erdfelder et al., 1996); between-subjects ANOVA, effect size f=0.43, α=0.05, power=0.80) resulted in an estimated 19 subjects per group. We therefore collected 25 full datasets per group (see below for participant exclusions) to ensure adequate power in the event of participant attrition. All participants were eligible for MR measurements and TMS interventions. They were free of medical, neurological, psychological or psychiatric conditions (including anxiety and depression as assessed by Beck Anxiety Inventory (Beck et al., 1988) and Beck Depression Inventory (Beck et al., 1961)) and were not taking any psychoactive or sleep-influencing medications at the time of the experiment. Participants showed no indications of abnormal sleep (as assessed with the Pittsburgh Sleep Quality Index (Buysse et al., 1989)) or excessive daytime sleepiness (as assessed with the Epworth sleepiness scale (Johns, 1991)) and were not considered extreme morning or evening types (as quantified with the Horne & Ostberg chronotype questionnaire (Horne and Ostberg, 1976)). Participants reported no previous extensive training with a musical instrument requiring dexterous finger movements (e.g., piano, guitar) or as a professional typist. None of the participants worked nights shifts or performed trans-meridian trips within the month prior to the experiment. Of these 75 complete datasets, 6 were excluded from the final analyses. As per our preregistration, participants were excluded from the analyses if they (1) failed to learn the sequence (N=0), (2) failed to perform the correct motor sequence (N=3 [2 control, 1 cTBS], as they presented performance accuracy >3SD below the mean of the sample), (3) did not comply with experimental instructions (N=3 [2 control, 1 iTBS], as they did not respect the regular sleep schedule), (4) presented movements in an fMRI session that exceed 2 voxels (N=0, but see below for 2 truncated scanning sessions due to movement), (5) showed anatomical abnormalities (N=0). Characteristics of the 69 participants included in the analyses are presented in Supplemental Table S7.

### 5.3 General experimental procedure

Participants first visited the MR unit for a baseline session including resting-state (RS), anatomical MRI measurements and baseline TMS measures (search of hotspot, resting motor threshold (rMT) and active motor threshold (aMT)). Note that RS data are not reported in the current study. At least three days after the baseline session, participants were invited for two consecutive experimental days (day 1 and day 2, see Figure 1). For the three days before experimental day 1 and until the end of the study (four nights in total), participants were instructed to follow a regular sleep/wake schedule (according to their own schedule ±1h, no naps). They were also instructed to refrain from alcohol and nicotine during this period. Sleep diaries and wrist actigraphy (ActiGraph wGT3X-BT, Pensacola, FL) were used to assess compliance to the sleep schedule.

On experimental day 1, participants were divided in 3 groups according to whether they received continuous, intermittent or control theta-burst stimulation (TBS) of the DLPFC (cTBS, iTBS or control, as described below; see Figure 1). TBS was applied outside the scanner before initial MSL (start experimental session between 9:30am and 5:30pm). Motor evoked potentials (MEPs; see Supplemental Methods and Results) were measured before and after TBS in order to probe corticospinal excitability. Immediately following stimulation, general motor execution (GME in Figure 1) was measured with a random serial reaction time task (see Supplemental Methods for a description of the task and Supplemental Results) and participants were then trained on the MSL task (as described below) while brain activity was recorded with fMRI (task duration: 17.17±3.96min, range 10.03-31.87min). The time between TBS offset and the start of MSL training (21.49±1.71min; range 16.33-25.83min) did not differ between groups (F_(2,66)_=.9 67, η_p_^2^=.296, *p*=.385). At the end of the experimental session on day 1, participants went home with the instructions to have a good night of sleep according to their sleep schedule, to not practice the task or consume any alcohol or drugs. Participants came back the next day (day 2) for a 24h task retest (start of MSL retest between 9:15am and 5:15pm) that took place in the MR scanner approximately at the same time as the training session on day 1 (time between start of MSL task on day 1 and day 2: 23.88±.88h, range 21.95-25.73h).

Prior to each MSL session on both experimental days, the psychomotor vigilance task (PVT; (Dinges and Powell, 1985)) and Stanford Sleepiness Scale (SSS; (Maclean et al., 1992)) were administered to provide objective and subjective assessments of vigilance, respectively (see Figure 1). Additionally, participants completed the St. Mary’s Hospital Sleep Questionnaire (Ellis et al., 1981) prior to each experimental session to report on the sleep quality of the preceding night. Sleep/vigilance scores prior to and during the study are reported in Supplemental Table S8.

### 5.4 Motor sequence learning task

Participants performed a bimanual finger-tapping task previously used in our group (Dolfen et al., 2019, 2021a), and implemented in Matlab. They practiced the task in an MRI scanner during two different sessions, i.e., MSL training and retest. During the task, participants tapped an eight-element finger sequence (8 fingers, all fingers except thumbs) as quickly and correctly as possible on a specially designed keyboard. Before initial training, participants were explicitly taught the sequence (4-7-3-8-6-2-5-1, with 1 representing the left little finger and 8 representing the right little finger). To do so, a short pre-training, consisting of slow and repeated sequence practice, was incorporated before each session (MSL training and MSL retest). Pre-training ended when three consecutive correct sequences were performed. Each session (MSL training and MSL retest) included 20 practice blocks followed, for the training session, by a 2min break and a post-training test including 4 practice blocks. This immediate post-training test was implemented to reduce the influence of fatigue on end-of-training performance (Pan and Rickard, 2015). During each practice block, the cross in the middle of the screen was green and the sequence of numbers corresponding to the fingers to press was displayed above the cross (see Figure 4A). Each practice block consisted of 48 keypresses (representing 6 correct sequences in an ideal practice block). After 48 keys were recorded, a 15s rest block was offered to the participants. During rest, the cross in the middle of the screen turned red and 8 asterisks were displayed instead of the sequence of numbers in order to avoid exposure to the sequence during rest (see Figure 4A). Participants were asked to look at the red fixation cross and to not move their fingers during rest blocks.

During MSL task performance, the timing and number of key presses were recorded, and performance was measured in terms of speed (mean time to perform a correct transition per block) and accuracy (percentage of correct transitions per block). Behavioral data were analyzed with separate repeated measures ANOVAs conducted on performance speed as well as accuracy during MSL training (20 blocks), immediate post-training test (4 blocks) as well as retest (20 blocks) with blocks as within-subject factor and group as between-subject factor (cTBS/iTBS/control).

Additionally, we performed exploratory analyses to examine consolidation processes on a micro timescale during initial MSL with similar procedures as in earlier work (Du et al., 2017; Bönstrup et al., 2019; Jacobacci et al., 2020; Buch et al., 2021). In line with these previous studies, the analysis at the micro timescale focused on performance speed. Specifically, micro-online gains in performance, reflecting performance changes during task practice periods, were defined as the difference between the average performance speed of the first 8 correct transitions of a block and the average speed of the last 8 correct transitions of the same block. The micro-online gains in performance extracted from the 20 practice blocks were then summed up for each participant for further statistical analyses. Micro-offline gains in performance, reflecting processes occurring during inter-practice rest periods, were computed as the difference between the average speed of the last 8 correct transitions of a block and the average speed of the first 8 correct transitions of the following block. The micro-offline gains in performance extracted from the 19 pairs of practice blocks were then summed up for each participant. ANOVAs were conducted for micro-online and -offline gains with group as between-subject factor (cTBS/iTBS/control). We then followed-up with two-tailed two-sample *t* tests to compare all pairs of groups. Based on data inspection and follow-up testing showing similar effects in the two active stimulation groups, we also compared performance between control and active stimulation groups (collapsed across cTBS and iTBS groups).

Last, we examined the effect of TMS on the behavioral markers of sleep-related consolidation occurring at the macro timescale. To do so, macro-offline gains in performance were computed as the percentage# (see Supplemental Table S6.1) of the difference in performance from the end of MSL training (average 4 blocks immediate post-training test) to the beginning of the MSL retest (average first 4 blocks) for both performance speed and accuracy. A three-way ANOVA with the factor group as between-subject factor (cTBS/iTBS/control) was conducted on macro-offline changes in performance speed and accuracy.

### 5.5 TMS administration

TMS was applied with a theta-burst stimulation (TBS) procedure (a burst of 3 pulses given at 50Hz, repeated every 200ms; (Huang et al., 2005)) on the DLPFC MNI coordinate −30 22 48 (Gann et al., 2021a) using a DuoMAG XT-100 rTMS stimulator (DEYMED Diagnostics s.r.o., Hronov, Czech Republic). The coil position was monitored online using neuronavigation (BrainSight, Rogue Research Inc, Montreal, Quebec, CA). To do so, individual T1 images were normalized in SPM8 and transferred into the BrainSight software with the corresponding normalization matrix to MNI space. Stimulation targets were created with the above-mentioned coordinate and a 45° angle so that the handle of the 70mm butterfly coil pointed posteriorly. This target was chosen as it has previously been shown to be functionally connected to the striatum and the hippocampus during rest and its stimulation was described to influence striatal as well as hippocampo-frontal functional connectivity during MSL (Gann et al., 2021a).

We applied intermittent (iTBS, 2s TBS trains repeated every 10s for 190s, 600 pulses) or continuous stimulation (cTBS, 40s uninterrupted train of TBS, 600 pulses) previously described to increase and decrease cortical excitability over the primary motor cortex (M1), respectively (Huang et al., 2005). Critically, the modulation of DLPFC cortical excitability is thought to be influenced by TBS similarly as in M1 (Tupak et al., 2013; Chung et al., 2017, 2018b). Active cTBS and iTBS were administered at 80% of the aMT (Huang et al., 2005). Control stimulation was applied with similar procedures as above but with a lower threshold (i.e., 40% aMT; (Bestmann et al., 2008; Heinen et al., 2011; van Nuenen et al., 2012; Romero et al., 2019; van Polanen et al., 2020)). This control stimulation procedure was chosen instead of e.g., sham stimulation approaches - such as tilting the coil away from the skull or adding an object between the coil and the head - in order to preserve similar sensory stimulation as during active stimulation. Active TBS effects (inhibitory and facilitatory) have been described to outlast the stimulation itself for up to 60min (Huang et al., 2005; Wischnewski and Schutter, 2015) and therefore overlapped with MSL training (MSL training ended on average 38.63±4.78min, range 29.23-53.33min, after the end of the TBS application).

### 5.6 Statistical Analyses of non-imaging data

Statistical analyses of the behavioral data (MSL, SRTT, PVT) as well as the MEP, sleep, questionnaire and demographic data were performed in SPSS Statistics 27 (IBM), with probability levels set to *p*<.05. We applied Greenhouse-Geisser corrections if the sphericity assumption was violated. *T* test statistics for independent sample tests were computed with un-pooled variance and correction of the degrees of freedom in the case of non-equal variance across two groups.

### 5.7 fMRI data acquisition and analysis

#### 5.7.1 Acquisition

During the baseline session, high-resolution T1-weighted structural images were acquired with a MPRAGE sequence (TR/TE=9.6/4.6ms; voxel size=0.98×0.98×1.2mm^3^; field of view=250×250×228mm^3^; 190 coronal slices) for each participant. Additional brain images described in the Supplemental Information were acquired during baseline but not analyzed in the presented paper.

Task-related fMRI data were acquired using an ascending gradient EPI pulse sequence for T2*-weighted images (TR=2000ms; TE=29.8ms; multiband factor 2; flip angle=90°; 54 transverse slices; slice thickness=2.5mm; interslice gap=0.2mm; voxel size=2.5×2.5×2.5mm^3^; field of view=210×210×145.6mm^3^; matrix=84×82; training: 514.23±115.90, post-test: 94.49±48.44, retest: 383.54±88.85 dynamical scans) during each task run. Note that one participant (cTBS group) did not finish the 20 MSL training blocks within the allocated scanning time of 30min, leaving fMRI data of only 19 full practice blocks. Additionally, due to scanner failure, the retest session of one participant (iTBS group) was performed without fMRI measurements (and therefore not included in any imaging contrasts involving the retest session).

After the last task run of each session, field maps were acquired (TR=1500ms; TE=3.5ms; flip angle=90°; 42 transverse slices; slice thickness=3.75mm; interslice gap=0.2mm; voxel size=3.75×3.75×3.75mm^3^; field of view=240×240×157.5mm^3^; matrix=64×64).

#### 5.7.2 Univariate fMRI analyses

##### 5.7.2.1 Spatial Preprocessing

Task-based functional volumes of each participant were realigned to the first image of each session and then realigned to the mean functional image computed across sessions using rigid body transformations# (see Supplemental Table S6.2). The mean functional image was co-registered to the high-resolution T1-weighted anatomical image using a rigid body transformation optimized to maximize the normalized mutual information between the two images. The resulting co-registration parameters were then applied to the realigned functional images. The structural image was segmented into gray matter, white matter, cerebrospinal fluid, bone, soft tissue, and background. We created an average subject-based template using DARTEL in SPM12, registered to the Montreal Neurological Institute (MNI) space. All functional and anatomical images were then normalized to the resulting template. Functional images were spatially smoothed using an isotropic 8mm full-width at half-maximum (FWHM) Gaussian kernel.

None of the participants were excluded after preprocessing as movement was overall minimal during scanning. The average ± SD translation and rotation across axes was 1.52±0.85mm and 1.47±0.78° (maximum absolute movement in translation=4.96mm and in rotation=4.05°) for MSL training, 0.62±0.61mm and 0.61±0.46° (maximum absolute movement in translation= 4.44mm and in rotation=2.55°) for MSL post-test and 1.22±0.79mm and 1.36±0.99° (maximum absolute movement in translation=4.27mm and in rotation=5.54°) for MSL retest. Movement parameters did not differ between groups (translation, TR: F_(2,66)_=0.558, η_p_^2^=0.017, *p*=0.575, PT: F_(2,66)_=0.132, η_p_^2^=0.004, *p*=0.876, RT: F_(2,65)_=1.235, η_p_^2^=0.037, *p*=0.298; rotation, TR: F_(2,66)_=2.204, η_p_^2^=0.063, *p*=0.118, PT: F_(2,66)_=2.334, η_p_^2^=0.066, *p*=0.105, RT: F_(2,65)_=0.766, η_p_^2^=0.023, *p*=0.469). However, MSL training data of 2 participants (1 cTBS, 1 iTBS) were truncated due to movement exceeding 2 voxels (i.e., the last 145 and last 56 scans - i.e., 33.33 and 10.31% of total scans, respectively - were excluded, leaving 13 and 18 full MSL practice blocks for the respective participants).

##### 5.7.2.2 Activation analyses

The pre-registered analysis of task-based fMRI data was based on a summary statistics approach and was conducted in 2 serial steps accounting for intra-individual (fixed effects) and inter-individual (random effects) variance, respectively. Changes in brain regional responses were estimated for each participant with a model including responses to the motor task and its linear modulation by performance speed (mean time to perform correct transitions per block; results presented in the supplements) for each task run (training (TR), immediate post-test (PT) and 24h-retest (RT)). The 15s rest blocks occurring between each block of motor practice served as the baseline condition modeled implicitly in the block design. These regressors consisted of box cars convolved with the canonical hemodynamic response function. Movement parameters derived from realignment as well as erroneous key presses were included as covariates of no interest. High-pass filtering was implemented in the design matrix using a cutoff period of 128s to remove slow drifts from the time series. Serial correlations in the fMRI signal were estimated using an autoregressive (order 1) plus white noise model and a restricted maximum likelihood (ReML) algorithm. Linear contrasts were generated at the individual level to test for the main effect of practice and its linear modulation by performance in each task run (TR and RT) as well as between task runs (TR vs. RT) # (see Supplemental Table S6.3). The resulting contrast images were further spatially smoothed (Gaussian kernel 6mm FWHM) and entered in a second level analysis for statistical inference at the group level (ANOVA with group (cTBS/iTBS/control) as between-subject factor), corresponding to a random effects model accounting for inter-subject variance. Follow-up analyses (two-sample *t* tests) on all pairs of groups are reported in the Supplemental Information, and exploratory analyses (two-sample *t* tests) comparing active (cTBS and iTBS) with control stimulation were performed when appropriate. Additionally, Supplemental Table S9 presents main effect of task practice across all experimental groups.

##### 5.7.2.3 Functional connectivity analyses

Methods and results corresponding to the pre-registered functional connectivity analyses are presented in the Supplemental Information.

##### 5.7.2.4 Regression analyses

Methods and results corresponding to the pre-registered regression analyses are presented in the Supplemental Information. In an additional exploratory analysis, we regressed the individuals’ contrast images from the activation-based analyses against the individuals’ micro-offline performance gains (speed) in a separate second level analysis for statistical inference at the group level (ANOVA with group (cTBS/iTBS/control) as between-subject factor), corresponding to a random effects model accounting for inter-subject variance. Follow-up analyses (*t* tests) on all pairs of groups and comparisons between active (cTBS and iTBS) and control stimulation were performed when appropriate.

##### 5.7.2.5 Statistical inferences

The set of voxel values resulting from each second level analysis listed above constituted maps of the F statistic testing for the main effect of group [SPM(F)] thresholded at *p*<0.005 (uncorrected for multiple comparisons). Follow-up two-sample *t* tests between all group pairs and between active (iTBS+cTBS) vs. control stimulation constituted maps of the T statistics [SPM(T)] and are reported in the main or the supplemental text. We used an ROI approach that included the DLPFC TBS target defined as a 10mm radius sphere around the target coordinate (−30 22 48mm) as well as the hippocampi and the basal ganglia (putamen, caudate nucleus and globus pallidus) defined with anatomical masks provided by Neuromorphometrics, Inc. (http://Neuromorphometrics.com/) under academic subscription and incorporated in SPM12. Statistical inference was conducted at a threshold of *p*<0.05 after family-wise error (FWE) correction for multiple comparisons over small volume within the ROIs (small volume correction (SVC) approach; (Poldrack, 2007; Poldrack et al., 2008)). SVC was applied on the number of voxels included in 10mm-radius spheres centered on coordinates from the literature (see Supplemental Table S10 for the main results and Supplemental Table S11 for the Supplemental Results). This procedure was followed by Holm-Bonferroni correction (Holm, 1979) for multiple brain regions highlighted in each contrast (*p*<0.05, indicated by an asterisk in the tables).

#### 5.7.3 Multivariate fMRI analyses

To investigate the effect of prefrontal stimulation on the neural processes supporting consolidation on a micro timescale, we performed exploratory multivariate analyses of the fMRI data of the initial MSL session. The goal of these exploratory analyses was to further characterize the stimulation-induced modulation of hippocampal and caudate activity during inter-practice rest periods (i.e., micro-offline epochs). Specifically, we investigated whether multivariate brain patterns observed during task practice in these brain regions persisted into the inter-practice rest periods during initial training. To do so, we computed the level of similarity (similarity index) of multi-voxel correlation structures (MVCS) between task practice and inter-practice rest periods in our two ROIs and in a control region. The analysis pipeline, implemented in Matlab, is described below and followed similar procedures as in our previous work (Gann et al., 2021b; King et al., 2021).

##### 5.7.3.1 Preprocessing

For each participant, the structural image was reoriented and segmented into gray matter (GM), white matter (WM), cerebrospinal fluid (CSF), bone, soft tissue, and background. The functional volumes were slice-time corrected (reference: middle slice) and then realigned to the mean functional image using rigid body transformations. This mean functional image was co-registered to the T1-weighted anatomical image using a rigid body transformation optimized to maximize the normalized mutual information between the two images. The resulting co-registration parameters were then applied to the realigned functional images. To optimize voxel pattern analyses, functional and anatomical data remained in subject-specific (i.e., native) space, and no spatial smoothing was applied to functional images (Tambini and Davachi, 2013; Gann et al., 2021b; King et al., 2021). Additional preprocessing of the time series was performed prior to running the MVCS analyses. Specifically, the whole-brain signal was detrended and high-pass filtered (cutoff=1/128). Framewise displacement of any given volume exceeding 0.5mm led to exclusion of that volume as well as the subsequent one (on average 8.65% of volumes excluded). Voxels with <10% GM probability were excluded from the analyses at the ROI level. To remove nuisance factors in each ROI, regression analyses were performed on the fMRI time-series of the remaining voxels. Specifically, regressors included the three first principal components of the signal extracted from the WM and CSF masks created during segmentation of the anatomical image (6 regressors) and the 6-dimensional head motion realignment parameters, as well as the realignment parameters squared, their derivatives, and the squared derivatives (24 regressors). Lastly, the number of volumes was matched between each task practice block and the following inter-practice rest block. To do so, we selected the x volumes around the middle volume of the longer block (practice or rest block), with x defined as the number of volumes in the block with the smaller number of scans (mean ± SD number of volumes per block state (practice or inter-practice rest) included in the analyses per group were for control: 122.24±18.45, cTBS: 115.38±20.71, iTBS: 118.67±20.16; the amount of volumes did not differ between the groups: F_(2,66)_=0.77, η_p_^2^=0.023, *p*=0.467).

##### 5.7.3.2 ROI selection and definition

The selection of ROIs for the MVCS analyses was based on the results of the univariate fMRI analyses showing stimulation-induced modulation of activity during inter-practice rest periods in the hippocampus and the caudate nucleus (see Table 1.1). Analyses also included a control ROI that did not show any stimulation-induced modulation of activity in the univariate analyses, even at a more permissive threshold. As signal to noise ratio and therefore similarity indices are usually lower in subcortical as compared to cortical regions (Gann et al., 2021b; King et al., 2021), we opted to select a *subcortical* control region, i.e., the thalamus, to facilitate qualitative comparisons between ROIs. Bilateral caudate, hippocampus and thalamus ROIs were therefore created in the native space of each individual using the FMRIB’s Integrated Registration Segmentation Toolkit (FSL FIRST; http://fsl.fmrib.ox.ac.uk/fsl/fslwiki/FIRST) employing boundary correction (‘fast’).

##### 5.7.3.3 Multivoxel correlation structure (MVCS) analyses

For each ROI (bilateral caudate, hippocampus and thalamus) and each block state (practice or rest), multi-voxel correlation structure (MVCS) matrices were computed with similar procedures as in previous research (Tambini and Davachi, 2013; Hermans et al., 2017; Gann et al., 2021b; King et al., 2021; Liu et al., 2021). Specifically, Pearson’s correlations were computed between each of n BOLD-fMRI voxel time courses, yielding an n by n MVCS matrix per ROI and per state. Pearson’s correlation coefficients were then Fisher Z-transformed to ensure normality. A similarity index (SI) reflecting the similarity of the multi-voxel patterns between the two block states (i.e., task practice and inter-practice rest) was computed as the r-to-z transformed correlation between the two MVCS matrices (Tambini and Davachi, 2013). SI between task and inter-practice rest volumes reflects the amount of persistence of task-related brain patterns into inter-practice rest periods in each experimental group (control, cTBS, iTBS). SI values were compared between groups using an ANOVA with group (cTBS/iTBS/control) as between-subject factor. Follow-up *t* tests on all pairs of groups and comparing active (cTBS and iTBS) to control stimulation were performed. All *t* tests including the control group were one-sided as we expected, based on the univariate fMRI results, less persistence of task patterns into rest after active (cTBS and iTBS) stimulation as compared to control stimulation.

Additionally, in order to test for relationships between behavioral and brain markers of micro-offline processes, we performed exploratory correlation analyses in SPSS between micro-offline gains in performance and (1) the amplitude of the BOLD response in ROIs reported in Table 1 (i.e., using parameter estimates extracted from the pre-registered univariate analyses reported in Table 1, Figure 3); and (2) SI in all ROIs. Correlations were compared between groups (cTBS/iTBS/control as well as active vs control) using an online tool available on http://home.ubalt.edu/ntsbarsh/business-stat/otherapplets/MultiCorr.htm.

## Author contributions

**Mareike Gann**: Conceptualization; Resources; Data curation; Software; Formal analysis; Investigation; Visualization; Writing - original draft; Writing - review and editing **Nina Dolfen**: Investigation; Methodology; Writing - review and editing **Bradley King**: Conceptualization; Resources; Data curation; Software; Formal analysis; Validation; Writing - review and editing **Edwin Robertson**: Conceptualization; Writing - review and editing **Geneviève Albouy**: Conceptualization; Resources; Data curation; Software; Formal analysis; Supervision; Funding acquisition; Validation; Investigation; Writing - original draft; Project administration; Writing - review and editing

## Data Availability

Data will be made available upon publication.

## Declaration of Competing Interest

The authors have no conflict of interest to declare.

## Acknowledgements

This work was supported by the Belgian Research Foundation Flanders (FWO; G099516N) and internal funds from KU Leuven. GA also received support from FWO (G0D7918N, G0B1419N, 1524218N) and Excellence of Science (EOS, 30446199, MEMODYN). MAG and ND received salary support from these grants. MAG is funded by a predoctoral fellowship from FWO (1141320N). Financial support for author BRK was provided by the European Union’s Horizon 2020 research and innovation program under the Marie Skłodowska-Curie grant agreement (703490) and a postdoctoral fellowship from FWO (132635). EMR received salary support from the Air Force Office of Scientific Research (AFOSR, Virginia, USA; FA9550-16-1-0191). We wish to thank Menno Veldman, Serena Reverberi, Simon Titone and Judith Nicolas as well as all involved students for assistance with data collection.

## Notes

### Competing Interest Statement

The authors have declared no competing interest.

