## Supplemental Information for "Prefrontal stimulation disrupts motor memory consolidation at the micro timescale"

### 1 Supplemental Methods

#### 1.1 General motor execution

General motor execution was measured after the stimulation but before MSL training using a random serial reaction time task (SRTT; (Nissen and Bullemer, 1987)), implemented in Matlab. This task was also performed inside the scanner but without acquiring the corresponding fMRI data. Participants saw eight squares on the screen, each representing one of the eight keys of the specifically designed keyboard and one of the 8 fingers (except thumbs). Green outlines of the squares indicated practice and red outlines rest blocks. For each green filled square appearing on the screen during the practice blocks, the key corresponding to the square's location had to be pressed as rapidly as possible. The following square was filled green after a key press (response-stimulus interval=0ms) according to a pseudorandom order and independent of the correctness of the previous key press. After 48 key presses the squares automatically turned red, indicating a rest block. During rest blocks (duration: 10s), participants were asked to look at the screen and to not move their fingers. The task included four practice blocks. During the performance of the SRTT, the timing and number of key presses was documented, and performance was measured in terms of speed (mean time to perform a correct key press per block) and accuracy (percentage of correct key presses per block). Repeated measures ANOVAs were conducted for speed as well as accuracy measures with blocks (4) as within-subject factor and group as between-subject factor (cTBS/iTBS/control), and the corresponding results are reported below.

#### 1.2 Motor evoked potentials

Motor evoked potentials (MEPs) were measured with a belly-tendon EMG montage on the right flexor dorsal interosseous (FDI) muscle. Active motor threshold (aMT) was characterized during voluntary submaximal FDI contraction as the lowest intensity for which minimum 5/10 MEPs were distinguishable from background EMG (Tambini et al., 2018; van Polanen et al., 2020). Resting motor threshold (rMT) was defined with single pulse stimulation of the M1 hotspot as the lowest intensity at which at least 5/10 MEPs measured on the FDI were larger than 50µV. As readout of corticospinal excitability changes in M1, twenty-one MEPs at 120% rMT were measured at two timepoints, i.e., pre- and post-TBS. Due to technical issues, MEP measurements are missing for 2 participants (1 control, 1 cTBS). Additionally, due to experimental error or interruption of the stimulation procedure because of participants' discomfort, MEP data of 7 participants (2 control, 4 cTBS, 1 iTBS) include less than 21 MEPs (with a minimum of 16 MEPs). The first MEP of each timepoint was excluded as their amplitudes are usually higher than subsequent MEPs due to reflex or startle responses. MEPs smaller than 50µV were excluded from the analysis (11.07%) and all remaining MEPs were visually inspected, leading to additional exclusions of 17 MEPs (0.72%)[#] (see Supplemental Table S6.4). Additionally, we performed an outlier analysis (3SD) on MEPs pre-TBS per participant, leading to no additional excluded MEPs. One participant was excluded from the MEP analyses due to background EMG noise (control group) and 6 additional participants (1 control, 2 cTBS, 3 iTBS) were excluded as they did not have enough remaining MEPs after the above-described exclusions (<10 MEPs left for pre- and/or post-TBS MEPs). MEPs of the remaining 60 participants (18 control, 21 cTBS, 21 iTBS) were averaged per timepoint and per individual (pre-TBS and post-TBS on experimental day 1, see Supplemental Figure S1) and entered into a repeated measures ANOVA with timepoint (pre-/ post-TBS) as within-subject factor and group (cTBS/iTBS/control) as between-subject factor. Additionally, changes in MEPs were calculated from pre- to post-TBS as the percentage[#] (see Supplemental Table S6.5) of difference between timepoints and correlated with offline gains in performance using Pearson's correlation. Percent change in MEPs were also used as covariates in fMRI regression analyses.

#### 1.3 Additional MR image acquisitions

During the baseline session we acquired additional T1 (TR/TE=8/3.8ms; voxel size=0.85×0.85×0.85mm$^3$; field of view=245×245×208.25mm$^3$; 245 sagittal slices, selective water excitation, bandwidth=299.3Hz) and T2 (TR/TE=2500/272ms; voxel size=0.85×0.85×0.85mm$^3$; field of view=245×245×190.4mm$^3$; 224 sagittal slices, bandwidth=876.8Hz) anatomical scans as well as RS fMRI data using an ascending gradient EPI pulse sequence for T2*-weighted images (TR=1000ms; TE=33ms; multiband factor 3; flip angle=80°; 42 transverse slices; interslice gap=0.5mm; voxel size=2.14×2.18×3mm$^3$; field of view=240×240×146.5mm$^3$; matrix=112×110; 300 dynamic scans) for each participant. During RS data acquisition, a dark screen (i.e., no visual stimuli) was presented and participants were instructed to remain still, close their eyes and to not think of anything in particular for the duration of the scan. The additional T1, T2 and RS data were not analyzed in the current study.

#### 1.4 Pre-registered fMRI analyses

##### 1.4.1 Functional connectivity analyses

Pre-registered psychophysiological interaction (PPI) analyses were computed to test task-related functional connectivity of the TBS target (-30 22 48mm) as well as additional seeds that were defined based on the results of the planned activation-based analyses (1 hippocampal and 2 caudate coordinates reported in Table 1.1). For each participant, experimental session and seed region of interest, the first eigenvariate was extracted using Singular Value Decomposition of the time series across the voxels included in a 10mm radius sphere centered around the seed of interest. A new linear model was generated at the individual level, using three regressors for each experimental task run. The first regressor corresponded to the BOLD activity in the seed region. The second regressor represented the practice of the learned sequence or the practice of the learned sequence modulated by performance speed. The third regressor represented the interaction of interest between the first (physiological) and the second (psychological) regressors. To build this regressor, the underlying neuronal activity were first estimated by a parametric empirical Bayes formulation, combined with the psychological factor, and subsequently convolved with the hemodynamic response function (Gitelman et al., 2003). The design matrix also included movement parameters as described for the activation-based analyses. Here, a significant PPI indicated a change in the regression coefficients (i.e., a change in the strength of the functional interaction) between any reported brain area and the seed region, related to the practice of the task or the performance speed changes during the practice of the task.

The resulting contrast images were further spatially smoothed (Gaussian kernel 6mm FWHM) and entered in a second level analysis for statistical inference at the group level (3-way ANOVA with group (cTBS/iTBS/control) as between-subject factor), corresponding to a random effects model accounting for inter-subject variance. The results related to functional connectivity can be found below.

##### 1.4.2 Regression analyses

As pre-registered, we regressed the individuals' contrast images from the activation-based analyses and the functional connectivity analyses against the individuals' macro-offline performance gains (speed and accuracy) and the individuals' change in MEPs in separate second level analyses for statistical inference at the group level (ANOVA with group (cTBS/iTBS/control) as between-subject factor), corresponding to a random effects model accounting for inter-subject variance. Follow-up analyses (*t* tests) on all pairs of groups and exploratory analyses (*t* tests) comparing active (cTBS and iTBS) with control stimulation were performed when appropriate. Results on planned regression analyses using macro-offline gains in speed and accuracy and changes in MEPs as covariates can be found below (Supplemental Tables S12-17).

### 2 Supplemental Results

#### 2.1 Pre-registered analyses

##### 2.1.1 Effect of prefrontal TBS on general motor execution

We investigated whether prefrontal stimulation altered general motor execution (GME) prior to initial training on experimental day 1 (see Figure 1). While performance improved over the course of the performed random serial reaction time task (speed: $F_{(2.116,139.684)}$=30.933, $\eta_p^2$=0.319, *p*<0.001; accuracy: $F_{(3, 198)}$=2.42, $\eta_p^2$=0.035, *p*=0.067), neither performance speed (main effect of group: $F_{(2,66)}$=0.352, $\eta_p^2$=0.011, *p*=0.704; group by block interaction: $F_{(4.233,139.684)}$=0.606, $\eta_p^2$=0.18, *p*=0.668) nor performance accuracy (main effect of group: $F_{(2,66)}$=2.225, $\eta_p^2$=0.063, *p*=0.116; group by block interaction: $F_{(6,198)}$=0.544, $\eta_p^2$=0.016, *p*=0.774) differed among groups.

##### 2.1.2 Effect of prefrontal TBS on MSL during test and retest

MSL test: After MSL training, participants were tested on the MSL task again (MSL test) after a short break allowing to reduce the influence of fatigue on end-of-training performance (Pan and Rickard, 2015). During the MSL test session, participants showed further performance speed improvement (main effect of block: $F_{(2.547,168.119)}$=4.035, $\eta_p^2$=0.058, $p$=0.012) and this effect was different between groups (block by group interaction: $F_{(5.095,168.119)}$=2.727, $\eta_p^2$=0.076, $p$=0.021). Follow-up analyses indicate that block-to-block performance changes differed between the cTBS and iTBS groups ($F_{(2.477,113.924)}$=5.039, $\eta_p^2$=0.099, $p$=0.005, $p_{Bonferroni}$=0.015) such that performance further improved across blocks in the iTBS group ($F_{(3,69)}$=4.68, $\eta_p^2$=0.169, $p$=0.005, $p_{Bonferroni}$=0.015) while it remained stable in the cTBS group ($F_{(1.771,40.731)}$=2.486, $\eta_p^2$=0.098, $p$=0.102). However, no significant main effect of group was observed for performance speed during MSL test ($F_{(2,66)}$=0.983, $\eta_p^2$=0.029, $p$=0.379). Performance accuracy remained stable during the test session (see Figure 2A, test panel: blocks 21-24; main effect of block: $F_{(2.57,169.638)}$=0.261, $\eta_p^2$=0.004, $p$=0.823) and this effect was similar in all groups (main effect of group: $F_{(2,66)}$=0.63, $\eta_p^2$=0.019, $p$=0.536; block by group interaction: $F_{(5.141,169.638)}$=0.13, $\eta_p^2$=0.021, $p$=0.619). In sum, these results indicate that performance speed reached more stable levels (i.e., a performance plateau) in the cTBS as compared to the iTBS group during the immediate post-training test.

MSL retest: During the retest session that took place approximately 24h (including a night of sleep) after initial learning, performance speed continued to improve during retest (main effect of block: $F_{(8.475,559.361)}$=23.68, $\eta_p^2$=0.264, $p$<0.001) and this effect was similar in all groups (main effect of group: $F_{(2,66)}$=0.889, $\eta_p^2$=0.026, $p$=0.416; block by group interaction: $F_{(16.95,559.361)}$=1.191, $\eta_p^2$=0.035, $p$=0.267). There were no further improvements in performance accuracy in any of the experimental groups (see Figure 2A, retest panel: blocks 25-44; main effect of block: $F_{(11.416,753.428)}$=1.546, $\eta_p^2$=0.023, $p$=0.107; main effect of group: $F_{(2,66)}$=0.485, $\eta_p^2$=0.014, $p$=0.618; block by group interaction: $F_{(22.831,753.428)}$=1.304, $\eta_p^2$=0.038, $p$=0.156).

Altogether, these behavioral results showed that performance speed reached more stable levels (i.e., a performance plateau) in the cTBS as compared to the iTBS group during the post-training test but this effect was no longer observed at the 24h retest.

##### 2.1.3 Effect of prefrontal TBS on corticospinal excitability

MEP values did not differ among groups ($F_{(1,57)}$=0.65, $\eta_p^2$=0.022, $p$=0.526) and showed no timepoint by group interaction ($F_{(2,57)}$=0.212, $\eta_p^2$=0.007, $p$=0.81). However, we observed a main effect of timepoint ($F_{(1,57)}$=13.217, $\eta_p^2$=0.188, $p$=0.001) driven by higher MEPs post-TBS compared to pre-TBS (see Supplemental Figure S1). Regression analyses between changes in MEPs amplitude and task-related activity/connectivity maps are reported in Supplemental Tables S16 and S17.

##### 2.1.4 Effect of prefrontal TBS on correlations between offline gains in performance and stimulation-induced changes in corticospinal excitability

We observed a significant correlation between offline gains in performance speed and MEP changes in the control group (r=0.491, $p$=0.039), but not in the cTBS (r=-0.237, $p$=0.3) or iTBS group (r=0.075, $p$=0.747). Differences[1] in correlations among groups were only trending ($\chi^2$=4.98, $p$=0.083).

we observed no significant correlation between offline gains in performance accuracy and MEP changes in the control (r=0.111, $p$=0.662), iTBS (r=-0.164, $p$=0.477) or cTBS groups (r=-0.038, $p$=0.869). Differences in correlations among groups were not significant ($\chi^2$=0.63, $p$=0.731).

##### 2.1.5 Effect of prefrontal TBS on dynamical brain activity during learning

We investigated whether prefrontal stimulation altered dynamical brain activity during learning, i.e., changes in brain activity as a function of block-to-block performance improvement. Parametric modulation analyses did not reveal any main effect of group during MSL training, retest or between sessions.

[1] Online tool used for calculations of differences between groups in correlations: http://home.ubalt.edu/ntsbarsh/business-stat/otherapplets/MultiCorr.htm

##### 2.1.6 Effect of prefrontal TBS on the relationship between task-related activation and macro-offline gains in performance speed

No group effects were observed in the regression analyses testing for relationships between macro-offline gains in performance and brain activity during training or changes in brain activity between training and retest sessions. However, we observed a significant group effect in the hippocampus in the regression map between task-related activation during MSL retest and macro-offline gains in performance speed (Supplemental Table S12.1). Specifically, higher hippocampal activity during inter-practice rest intervals at retest was related to greater macro-offline gains in performance in the control group as compared to the active TBS groups (see Supplemental Figure S2A, Supplemental Table S13).

Additionally, the relationship between dynamical brain activity (as assessed with parametric modulations) and macro-offline gains in performance speed differed between groups in the caudate, putamen and globus pallidus during the retest session (Supplemental Table S12.2). Specifically, macro-offline gains in performance were associated to a progressive decrease in basal ganglia activity in the control group while it was related to a practice-related increase in the iTBS group (see Supplemental Figure S2B and Supplemental Table S13 for paired comparisons showing significant differences between control and the two active stimulation groups as well as between iTBS and cTBS groups).

Similarly, inter-session changes in dynamical activity of the hippocampus, caudate nucleus, putamen and globus pallidus were differently related to macro-offline gains in performance speed between groups (Supplemental Table S12.2). Inspection of dynamical patterns in the basal ganglia indicated that poorer macro-offline gains in performance were related to higher learning-related modulation of brain activity at retest as compared to training in the control group while an opposite pattern of results was observed in the iTBS group (see Supplemental Table S13 for paired comparisons showing significant differences between control and the two active stimulation groups as well as between iTBS and cTBS groups). In contrast, in the hippocampus, higher macro-offline gains in performance were related to the change in dynamics between training and retest (i.e., decrease as function of practice during training and increase in proportion to performance improvement during retest) that was more pronounced in the control group as compared to the cTBS group (Supplemental Table S13).

Altogether, these results suggest that prefrontal stimulation modulated the relationship between macro-offline gains in performance speed and activity in the hippocampus and the striatum. Specifically, active stimulation interrupted the link - observed under control conditions – between rest- and task-related hippocampal activity and macro-offline gains in performance. Similar effects were observed in the striatum, and they were further accentuated under iTBS whereby the brain-behavior relationship was flipped as compared to controls.

###### 2.1.7 Effect of prefrontal TBS on task-related functional connectivity

As per our pre-registration, we investigated whether the functional connectivity of the seed regions highlighted in the activation-based analyses (Table 1.1) and of the DLPFC TMS target differed among groups. There were no significant group effects on connectivity maps in the ROIs.

###### 2.1.8 Effect of prefrontal TBS on the relationship between task-related functional connectivity and macro-offline gains in performance speed

We observed that the relationship between macro-offline gains in performance speed and dynamical connectivity (as assessed with parametric modulations) between the DLPFC and the caudate nucleus as well as the globus pallidus differed between groups during initial MSL training (Supplemental Table S14.1). Specifically, larger practice-related decreases in DLPFC-caudate connectivity were related to greater overnight gains in performance speed in the control group while such relationships were not observed in the active stimulation groups (see Supplemental Figure S2C, Supplemental Table S15).

Altogether, these results suggest that active stimulation disturbed the relationships between macro-offline gains in performance speed and dynamical connectivity patterns of the DLPFC during initial training.

### 3 Supplemental Tables

**Supplemental Table S1.** Behavioral results of processes at the micro timescale – pairwise group comparisons (*t* tests)

| | cTBS vs. control | iTBS vs. control | cTBS vs. iTBS |
|---|---|---|---|
| **micro-offline gains** | $t_{(32.390)}$=-2.298, d=0.708, *p*=0.028 | $t_{(43)}$=-1.517, d=-0.453, *p*=0.137 | $t_{(46)}$=0-.781, d=0-.225, *p*=.439 |
| **micro-online gains** | $t_{(43)}$=2.348, d=0.702, *p*=0.024 | $t_{(43)}$=1.368, d=0.409, *p*=0.178 | $t_{(46)}$=0.973, d=0.281, *p*=0.336 |
| **overall learning** | $t_{(43)}$=0.160, d=0.048, *p*=0.873 | $t_{(43)}$=-0.718, d=-0.215, *p*=0.476 | $t_{(46)}$=0.855, d=0.247, *p*=0.397 |

**Supplemental Table S2.** Functional imaging results of activation-based contrasts – pairwise group comparisons (*t* tests)

| Area | x | y | z | k | Z | $p_{FWEsvc}$ |
|---|---|---|---|---|---|---|
| **Main effect of practice** | | | | | | |
| ***TRAINING*** | | | | | | |
| *[cTBS-control]* | | | | | | |
| Hippocampus | -30 | -10 | -26 | 48 | 3.1 | 0.021* |
| Caudate | -8 | 16 | 4 | 63 | 2.99 | 0.028 |
| *[iTBS-control]* | | | | | | |
| Hippocampus | -28 | -10 | -26 | 111 | 3.5 | 0.007* |
| Caudate | -10 | 16 | 4 | 4654 | 3.35 | 0.01* |
| *[control-cTBS] \| [control-iTBS] \| [cTBS-iTBS] \| [iTBS-cTBS]* | | | | | | |
| No significant responses in the ROIs | | | | | | |
| ***TRAINING - RETEST*** | | | | | | |
| *[cTBS-control]* | | | | | | |
| Caudate | -6 | 12 | 6 | 188 | 3.24 | 0.015* |
| *[iTBS-control]* | | | | | | |
| Caudate | -6 | 12 | 6 | 288 | 3.05 | 0.024* |
| *[control-cTBS] \| [control-iTBS] \| [cTBS-iTBS] \| [iTBS-cTBS]* | | | | | | |
| No significant responses in the ROIs | | | | | | |

*Asterisk (*) indicates significance at p<0.05 after Holm-Bonferroni correction for multiple comparison. Statistics were extracted from regions of interest (ROI) without applying anatomical masks.*

**Supplemental Table S3.** Functional imaging results of persistence of task-related brain patterns into inter-practice rest intervals – pairwise group comparisons (*t* tests)

| | control vs. cTBS (one-sided *t* test) | control vs. iTBS (one-sided *t* test) | cTBS vs. iTBS (2-sided *t* test) |
|---|---|---|---|
| **caudate nucleus** | $t_{(43)}$=1.582, d=-0.473, *p*=0.061 | $t_{(43)}$=1.734, d=-0.518, *p*=0.045 | $t_{(46)}$=0.255, d=-0.074, *p*=0.8 |
| **hippocampus** | $t_{(43)}$=1.352, d=-0.404, *p*=0.092 | $t_{(43)}$=1.266, d=-.378, *p*=.106 | $t_{(46)}$=0.021, d=-0.006, *p*=0.984 |
| **thalamus** | $t_{(43)}$=0.656, d=-0.196, *p*=0.258 | $t_{(43)}$=1.233, d=-.366, *p*=.114 | $t_{(46)}$=0.676, d=0.195, *p*=0.502 |

**Supplemental Table S4.** Group pair comparisons regarding the relationship between brain activity and pattern persistence as well as micro-offline gains and pattern persistence

| | cTBS | iTBS | active | control | cTBS vs. iTBS | cTBS vs. control | iTBS vs. control |
|---|---|---|---|---|---|---|---|
| | ***correlation between brain activity (see Table 1.1 of main text) and MVCS pattern persistence*** | | | | | | |
| **caudate nucleus** | r=-0.089; p=0.679 | r=-0.168; p=0.434 | r=-0.137; p=0.354 | r=-0.266; p=0.243 | $\chi^2$=0.07, p=0.795 | $\chi^2$=0.33, p=0.568 | $\chi^2$=0.10, p=0.749 |
| **hippocampus** | r=0.247; p=0.245 | r=-0.074; p=0.731 | r=0.049; p=0.74 | r=-0.56; p=0.008 | $\chi^2$=1.12, p=0.29 | $\chi^2$=7.59, p=0.006 | $\chi^2$=3.03, p=0.082 |
| | ***correlation between micro-offline gains and MVCS pattern persistence*** | | | | | | |
| **caudate nucleus** | r=-0.356; p=0.087 | r=0.108; p=0.614 | r=-0.081; p=0.585 | r=-0.063; p=0.787 | $\chi^2$=2.4, p=0.119 | $\chi^2$=0.93, p=0.336 | $\chi^2$=0.29, p=0.593 |
| **hippocampus** | r=-0.145; p=0.499 | r=-0.029; p=0.892 | r=-0.074; p=0.617 | r=-0.032; p=0.89 | $\chi^2$=0.143, p=0.705 | $\chi^2$=0.126, p=0.723 | $\chi^2$=0.00009, p=0.993 |
| **thalamus** | r=-0.225; p=0.291 | r=0.178; p=0.406 | r=0.013; p=0.932 | r=0.165; p=0.474 | $\chi^2$=1.76, p=0.185 | $\chi^2$=1.52, p=0.218 | $\chi^2$=0.002, p=0.965 |

*MVCS: multivoxel correlation structure.*

**Supplemental Table S5.** Functional imaging results of the regression analyses between task-related activity maps (main effect of practice) and the sum of micro-offline gains during initial MSL

| **Area** | **x** | **y** | **z** | **k** | **Z** | $p_{FWEsvc}$ |
|---|---|---|---|---|---|---|
| **Pairwise group comparisons (*t* tests)** | | | | | | |
| *[iTBS-control]* | | | | | | |
| Hippocampus | 26 | -30 | -6 | 3 | 2.77 | 0.048* |
| *[control-iTBS]\|[control-cTBS]\|[cTBS-control]\|[iTBS-cTBS]\|[cTBS-iTBS]* | | | | | | |
| No significant responses | | | | | | |

*Asterisk (*) indicates significance at p<0.05 after Holm-Bonferroni correction for multiple comparison.*

**Supplemental Table S6.** Deviations from preregistration

1. **Offline gains**

**Preregistration:** Offline gains in performance speed and accuracy will be computed as the difference in performance from the end of Training (average 4 blocks immediate post-Training test) to the beginning of the MSL Retest (average first 4 blocks).
**Manuscript:** Offline gains were computed as the percentage change from performance at the end of training (average 4 blocks immediate post-training test) to the beginning of the MSL retest (average first 4 blocks).
**Justification:** Baseline performance levels during post-training test being more variable than expected, we normalized the offline gains to the performance level during post-training test.

2. **Field maps**

**Preregistration:** Task-based functional volumes of each participant will be corrected for inhomogeneities in the magnetic field (i.e., unwarped) using the field map images.
**Manuscript:** Field maps were not used in the current preprocessing.
**Reason:** The introduction of the field maps in the SPM preprocessing pipeline induced unforeseen image distortions in some participants. We elected to not use the field maps in the final pipeline in order to stay consistent across individuals.

3. **Linear contrasts**

**Preregistration:** Linear contrasts will be generated at the individual level to test for the main effect of practice and its linear modulation by performance in each task run (TR, PT and RT) as well as between task runs (TR vs. PT, TR vs. RT, PT vs. RT).
**Manuscript:** Post-tests (PT) linear contrasts are not reported in the present study.

4. **Motor evoked potential (MEP) preprocessing**

**Preregistration:** Individual MEP trials will be excluded from analyses if the measured variable is greater than 3 standard deviations above or below the participant's mean.
**Manuscript:** Additional exclusions:

- Of 1$^{st}$ MEPs of each timepoint per participant.
- MEPs with amplitudes smaller than 50µV.

Based on visual inspection of shape and background noise.
**Justification:** Additional exclusion criteria were used as the MEP data were noisier than expected.

- 1$^{st}$ MEP: First MEPs often show higher amplitudes due to reflex/startle responses.
- <50µV: Potentials smaller than 50µV are not considered as MEPs.
- Visual inspection: Some MEPs were discarded if they did not show the usually observed waveform or if background EMG noise was present.

5. **MEP change**

**Preregistration:** Changes in MEPs will be calculated from pre- to post-TBS as the difference between time points.
**Manuscript:** Changes in MEPs were calculated as the percentage change from pre- to post-TBS.
**Justification:** Baseline MEP levels pre-TBS being more variable then expected (see Supplemental Figure S1), we normalized the MEP change to the MEP level pre-TBS.

*MEPs: motor evoked potentials, TBS: theta-burst stimulation, TR: training, PT: post-test, RT: retest.*

**Supplemental Table S7.** Participant characteristics

| | Control group | cTBS group | iTBS group | Main effect of group |
|---|---|---|---|---|
| N (female) | 21 (14) | 24 (16) | 24 (16) | $\chi^2_{(2)}$=0, *p*=1 |
| Age (years) | 23.19 ± 2.96 | 23.5 ± 2.43 | 23.58 ± 2.24 | $F_{(2,66)}$=0.15 , *p*=0.87 |
| Edinburgh Handedness | 86.67 ± 12.08 | 82.92 ± 16.28 | 84.79 ± 14.10 | $F_{(2,66)}$=0.38, *p*=0.68 |
| Epworth Sleepiness Scale | 6.33 ± 3.37 | 6.38 ± 3.41 | 7.29 ± 3.5 | $F_{(2,66)}$=0.59, *p*=0.56 |
| PSQI | 2.48±1.25 | 2.46±1.285 | 2.17±1.2 | $F_{(2,66)}$=0.46, *p*=0.64 |
| Chronoscore (CRQ)[a] | 48.95 ± 7.8 | 51.13 ± 6.17 | 51 ± 8.66 | $F_{(2,64)}$=0.55, *p*=0.58 |
| Beck Depression Scale | 3.14 ± 2.59 | 3.25 ± 3.6 | 2.17 ± 3.03 | $F_{(2,66)}$=0.86, *p*=0.43 |
| Beck Anxiety Scale | 2.62 ± 2.54 | 2.46 ± 1.64 | 2.83 ± 3.27 | $F_{(2,66)}$=0.13, *p*=0.88 |

*Values are means ± standard deviation. p values are based on one-way ANOVAs with the between-subject factor group (3). PSQI: Pittsburgh Sleep Quality Index, CRQ: Circadian Rhythm Questionnaire. [a]Chronoscores could not be determined for 2 participants (1 control, 1 iTBS) due to one missing answer each. None of the participants were categorized as extreme morning or evening types.*

**Supplemental Table S8.** Sleep/vigilance scores

| | Control | cTBS | iTBS | Statistical analyses |
|---|---|---|---|---|
| **Sleep duration (h)** | | | | Main effect of group: $F_{(2,66)}$=0.76, *p*=0.47 |
| Mean (4 nights) | 8.7 ± 1.08 | 8.46 ± .98 | 8.7 ± 1.05 | Main effect of night: $F_{(3,198)}$=1.93, *p*=0.13 |
| Night 1 | 9.05 ± 1.19 | 8.32 ± 1.03 | 8.52 ± 1.17 | Night x group interaction: $F_{(6198)}$=1.72, *p*=0.12 |
| Night 2 | 8.81 ± 1.12 | 8.52 ± 0.99 | 8.84 ± 0.97 | |
| Night 3 | 8.42 ± 1 | 8.36 ± 1 | 8.49 ± 0.97 | |
| Night 4 | 8.5 ± 0.97 | 8.64 ±0.91 | 8.94 ± 1.07 | |
| **St. Mary's Sleep quality** | | | | Main effect of group: $F_{(2,66)}$=0.31, *p*=0.73 |
| Night 3 | 4.14±0.66 | 3.92±0.83 | 4±0.59 | Main effect of night: $F_{(1,66)}$=1.07, *p*=0.31 |
| Night 4 | 4.14±0.36 | 4.13±0.85 | 4.13±0.68 | Night x group interaction: $F_{(2,66)}$=0.31, *p*=0.74 |
| **Psychomotor vigilance task (s)** | | | | Main effect of group: $F_{(2,66)}$=0.70, *p*=0.5 |
| Training | 0.29±0.02 | 0.28±0.02 | 0.28±0.03 | Main effect of session: $F_{(1,66)}$=2.297, *p*=0.134 |
| Retest | 0.28±0.02 | 0.28±0.02 | 0.28±0.03 | Session x group interaction: $F_{(2,66)}$=3.121, *p*=0.051 |
| **Stanford sleepiness score** | | | | Main effect of group: $F_{(2,64)}$=1.96, *p*=0.15 |
| Training | 1.93 ±0.68 | 1.71 ±0.61 | 1.85 ±0.8 | Main effect of session: $F_{(1,64)}$=0.13, *p*=0.72 |
| Retest | 2.08 ±0.66 | 1.65 ±0.68 | 1.65 ±0.57 | Session x group interaction: $F_{(2,64)}$=0.96, *p*=0.39 |

*Values are means ± standard deviation. p values are based on repeated measures ANOVAs with the between-subject factor group (3) and sessions/nights as within-subject factor. Sleep duration was assessed based on actiwatch and sleep diary data. Night 1 corresponds to the night 3 days before the experiment and Night 4 to the night between training and retest. Stanford sleepiness scores are missing for 2 participants (1 iTBS, 1 control).*

**Supplemental Table S9.** Functional imaging results for initial MSL training across groups

| Area | x | y | z | k | T | $p_{FWE}$ |
|---|---|---|---|---|---|---|
| **1. Main effect of practice** | | | | | | |
| ***+TRAINING*** | | | | | | |
| Sensorimotor cortex (extending to premotor cortex, insula, putamen) | -30 | -16 | 52 | 52774 | 21.49 | <0.001 |
| Inferior parietal | -44 | -34 | 40 | | 19.88 | <0.001 |
| Superior parietal | -32 | -50 | 56 | | 19.82 | <0.001 |
| Cerebellum | 20 | -54 | -24 | 18037 | 20.16 | <0.001 |
| | 26 | -64 | -54 | | 19.64 | <0.001 |
| | 18 | -66 | -48 | | 19.54 | <0.001 |
| Middle frontal | 38 | 38 | 26 | 66 | 5.71 | <0.001 |
| ***-TRAINING*** | | | | | | |
| Middle cingulum (extending to temporal areas, frontal areas, parietal areas, occipital areas, precuneus, parahippocampus, hippocampus, insula, caudate, putamen) | -4 | -32 | 40 | 79124 | 22.7 | <0.001 |
| Angular gyrus | -52 | -60 | 30 | | 19.37 | <0.001 |
| | -44 | -70 | 34 | | 19.1 | <0.001 |
| Cerebellum | 36 | -76 | -38 | 1123 | 12.84 | <0.001 |
| | -24 | -80 | -36 | 886 | 10.93 | <0.001 |
| | 2 | -58 | -50 | 55 | 7.11 | <0.001 |
| ***2.* Modulation in activity by performance speed** | | | | | | |
| ***Brain regions wherein activity increases with practice*** | | | | | | |
| M1 | 38 | -22 | 54 | 919 | 8.85 | <0.001 |
| | -36 | -24 | 56 | 454 | 7.07 | <0.001 |
| Cerebellum | -18 | -52 | -24 | 590 | 6.73 | <0.001 |
| | -4 | -60 | -16 | | 5.39 | <0.001 |
| | 20 | -52 | -26 | | 4.97 | <0.001 |
| Putamen | 22 | 8 | 8 | 742 | 6.62 | <0.001 |
| | 22 | 10 | -6 | | 5.85 | <0.001 |
| | -22 | 14 | 0 | 787 | 6.54 | <0.001 |
| Caudate | -14 | 6 | 18 | | 4.84 | |
| ***Brain regions wherein activity decreases with practice*** | | | | | | |
| No significant responses at $p_{FWE}$<0.05 | | | | | | |

*Significant results at $p_{FWE}$<0.05 (whole-brain family-wise error (FWE) corrected). M1: primary motor cortex.*

**Supplemental Table S10.** Coordinates of areas of interest used for spherical small volume corrections for the main results

| Area | x mm | y mm | z mm | Reference |
|---|---|---|---|---|
| ***PARA-/HIPPOCAMPAL*** | | | | |
| Posterior Hippocampus | ±24 | -34 | 2 | (Albouy et al., 2008) |
| Hippocampus | ±34 | -10 | -20 | (Albouy et al., 2015) |
| ***STRIATAL*** | | | | |
| Caudate | ±9 | 15 | -3 | (Schendan et al., 2003) |
| Caudate | ±15 | 12 | 12 | (Schendan et al., 2003) |
| Globus Pallidus | ±12 | 2 | 0 | (Lehéricy et al., 2006) |
| ***TMS target*** | | | | |
| DLPFC | -30 | 22 | 48 | (Gann et al., 2021) |

*Coordinates in MNI space. DLPFC: dorsolateral prefrontal cortex, MNI: Montreal Neurological Institute.*

**Supplemental Table S11.** Coordinates of areas of interest used for spherical small volume corrections for the supplemental results

| Area | x mm | y mm | z mm | Reference |
|---|---|---|---|---|
| ***PARA-/HIPPOCAMPAL*** | | | | |
| Hippocampus head/ Parahippocampal gyrus | ±33 | -18 | -27 | (Schendan et al., 2003) |
| Posterior Hippocampus | ±24 | -34 | 2 | (Albouy et al., 2008) |
| Hippocampus | ±34 | -10 | -20 | (Albouy et al., 2015) |
| Hippocampus | ±26 | -28 | -22 | (Albouy et al., 2008) |
| Hippocampus | ±22 | -28 | -10 | (Albouy et al., 2008) |
| Hippocampus | ±16 | -14 | -28 | (Albouy et al., 2008) |
| Hippocampus | ±32 | -28 | -10 | (Albouy et al., 2008) |
| ***STRIATAL*** | | | | |
| Caudate | ±9 | 15 | -3 | (Schendan et al., 2003) |
| Caudate | ±15 | 12 | 12 | (Schendan et al., 2003) |
| Caudate | ±15 | -6 | 18 | (Schendan et al., 2003) |
| Putamen | ±27 | -9 | 9 | (Schendan et al., 2003) |
| Putamen | ±24 | 4 | 16 | (Peigneux et al., 2000) |
| Putamen | ±18 | 0 | -8 | (van der Graaf et al., 2004) |
| Globus Pallidus | ±12 | 2 | 0 | (Lehéricy et al., 2006) |
| ***TMS target*** | | | | |
| DLPFC | -30 | 22 | 48 | (Gann et al., 2021) |

*Coordinates in MNI space. DLPFC: dorsolateral prefrontal cortex, MNI: Montreal Neurological Institute.*

**Supplemental Table S12.** Functional imaging results of the regression analyses between activity maps and overnight macro-offline gains in performance speed – main effect of group (F test)

| Area | x | y | z | k | Z | $p_{FWEsvc}$ |
|---|---|---|---|---|---|---|
| **1. Main effect of practice** | | | | | | |
| ***TRAINING*** | | | | | | |
| No significant responses | | | | | | |
| ***RETEST*** | | | | | | |
| Posterior Hippocampus | -26 | -40 | -2 | 8 | 3.12 | 0.024* |
| Anterior Hippocampus | -30 | -14 | -26 | 64 | 3.02 | 0.031 |
| | 26 | -8 | -22 | 23 | 2.88 | 0.044 |
| ***TRAINING – RETEST*** | | | | | | |
| No significant responses | | | | | | |
| ***2.* Modulation in activity by speed** | | | | | | |
| ***TRAINING*** | | | | | | |
| No significant responses | | | | | | |
| ***RETEST*** | | | | | | |
| Caudate | 8 | 10 | 4 | 86 | 3.16 | 0.023 |
| | -12 | 20 | -8 | 6 | 3.18 | 0.021 |
| | -6 | 10 | 4 | 24 | 3.03 | 0.032 |
| | -16 | 0 | 24 | 12 | 3.11 | 0.026 |
| Globus Pallidus | 18 | 0 | 4 | 237 | 3.43 | 0.010* |
| Putamen | 22 | 0 | 8 | | 3.07 | 0.029 |
| | -30 | -4 | 4 | 96 | 3.25 | 0.018 |
| ***TRAINING – RETEST*** | | | | | | |
| Caudate | -16 | 2 | 22 | 30 | 3.90 | 0.002* |
| Hippocampus | 36 | -28 | -10 | 11 | 3.24 | 0.018 |
| Globus Pallidus | 18 | 0 | 4 | 76 | 3.14 | 0.024 |
| Putamen | 24 | 18 | -6 | 13 | 2.93 | 0.042 |
| | -28 | -4 | 4 | 50 | 2.92 | 0.043 |

*Asterisk (*) indicates significance at p<0.05 after Holm-Bonferroni correction for multiple comparison. Statistics were extracted from regions of interest after applying anatomical masks.*

**Supplemental Table S13.** Functional imaging results of the regression analyses between activity maps and overnight macro-offline gains in performance speed – pairwise group comparisons (*t* tests)

| Area | x | y | z | k | Z | $p_{FWEsvc}$ |
|---|---|---|---|---|---|---|
| **1. Main effect of practice** | | | | | | |
| ***RETEST*** | | | | | | |
| *[control-cTBS]* | | | | | | |
| Caudate | 16 | 8 | 10 | 174 | 2.85 | 0.038* |
| | -14 | 0 | 24 | 88 | 2.82 | 0.04* |
| *[cTBS-control]* | | | | | | |
| Posterior Hippocampus | -24 | -42 | -4 | 89 | 3.33 | 0.01* |
| Anterior Hippocampus | -32 | -14 | -28 | 1582 | 3.56 | 0.005* |
| | 26 | -10 | -22 | 73 | 3 | 0.026 |
| *[control-iTBS]* | | | | | | |
| Caudate | -14 | 0 | 24 | 93 | 2.93 | 0.031* |
| *[iTBS-control]* | | | | | | |
| Posterior Hippocampus | -26 | -40 | -4 | 1142 | 3.89 | 0.002* |
| Anterior Hippocampus | -26 | -16 | -22 | 2185 | 3.57 | 0.005* |
| | 26 | -8 | -22 | 227 | 3.51 | 0.006* |
| ***2.* Modulation in activity by speed** | | | | | | |
| ***RETEST*** | | | | | | |
| *[control-cTBS]* | | | | | | |
| Putamen | 22 | -2 | 6 | 561 | 3.05 | 0.024* |
| *[cTBS-iTBS]* | | | | | | |
| Caudate | 10 | 6 | 6 | 6639 | 2.89 | 0.036 |
| Putamen | -28 | -4 | 6 | 181 | 3.39 | 0.009* |
| *[control-iTBS]* | | | | | | |
| Caudate | 14 | 6 | 4 | 4634 | 3.61 | 0.005* |
| Putamen | 20 | -2 | 6 | | 3.89 | 0.002* |
| | -34 | -4 | 4 | | 3.54 | 0.006* |
| ***TRAINING – RETEST*** | | | | | | |
| *[control-cTBS]* | | | | | | |
| Hippocampus | 36 | -28 | -10 | 54 | 3.62 | 0.005* |
| *[cTBS-control]* | | | | | | |
| Putamen | 18 | 0 | 2 | 51 | 2.84 | 0.041* |
| *[iTBS-cTBS]* | | | | | | |
| Putamen | -26 | 4 | 4 | 158 | 3.12 | 0.02* |
| Caudate | -8 | 12 | 6 | 2499 | 3.06 | 0.024* |
| *[iTBS-control]* | | | | | | |
| Putamen | -32 | -2 | 4 | 3858 | 3.27 | 0.013* |
| | 22 | -2 | 4 | | 3.61 | 0.005* |
| Caudate | 10 | 8 | 4 | | 3.11 | 0.021* |

*Asterisk (*) indicates significance at p<0.05 after Holm-Bonferroni correction for multiple comparison. Statistics were extracted from regions of interest without applying anatomical masks.*

**Supplemental Table S14.** Functional imaging results of the regression analyses between connectivity maps and overnight macro-offline gains in performance speed – main effect of group (F test)

| Area | x | y | z | k | Z | $p_{FWEsvc}$ |
|---|---|---|---|---|---|---|
| **1. TRAINING** | | | | | | |
| ***Hippocampus [-28 -10 -26] connectivity*** | | | | | | |
| No significant responses | | | | | | |
| ***Caudate [-8 16 4] connectivity*** | | | | | | |
| No significant responses | | | | | | |
| ***DLPFC target [-30 22 48] connectivity*** | | | | | | |
| No significant responses | | | | | | |
| ***DLPFC target [-30 22 48] modulation of connectivity by speed*** | | | | | | |
| Caudate | 14 | 20 | 6 | 119 | 4.14 | 0.001* |
| | -12 | 10 | -2 | 7 | 3.16 | 0.024* |
| Globus Pallidus | -14 | 4 | -2 | 81 | 3.87 | 0.003* |
| | 18 | -2 | -4 | 96 | 3.32 | 0.015* |
| **2. TRAINING – RETEST** | | | | | | |
| ***Caudate [-6 12 6] connectivity*** | | | | | | |
| No significant responses | | | | | | |

*Asterisk (*) indicates significance at p<.05 after Holm-Bonferroni correction for multiple comparison. Statistics were extracted from regions of interest after applying anatomical masks. DLPFC: dorsolateral prefrontal cortex.*

**Supplemental Table S15.** Functional imaging results of the regression analyses between connectivity maps and overnight macro-offline gains in performance speed – pairwise group comparisons (*t* tests)

| Area | x | y | z | k | Z | $p_{FWEsvc}$ |
|---|---|---|---|---|---|---|
| **TRAINING** | | | | | | |
| ***Modulation in DLPFC target [-30 22 48] connectivity by speed*** | | | | | | |
| *[control-cTBS]* | | | | | | |
| Caudate | 14 | 20 | 4 | 1181 | 4.49 | <0.001* |
| Putamen | 22 | -4 | 2 | | 2.8 | 0.047 |
| *[control-iTBS]* | | | | | | |
| Caudate | 14 | 20 | 4 | 3753 | 4.59 | <0.001* |
| Putamen | 18 | -8 | 6 | | 3.33 | 0.011* |

*Asterisk (*) indicates significance at p<0.05 after Holm-Bonferroni correction for multiple comparison. Statistics were extracted from regions of interest without applying anatomical masks.*

**Supplemental Table S16.** Functional imaging results of the regression analyses between activity maps and overnight macro-offline gains in accuracy as well as changes in MEPs – main effect of group (F test)

| Area | x | y | z | k | Z | $p_{FWEsvc}$ |
|---|---|---|---|---|---|---|
| **1. Main effect of practice: Regression with macro-offline gain in accuracy** | | | | | | |
| ***TRAINING \| RETEST \| TRAINING – RETEST*** | | | | | | |
| No significant responses | | | | | | |
| **2. Modulation in activity by speed: Regression with macro-offline gain in accuracy** | | | | | | |
| ***TRAINING \| RETEST \| TRAINING – RETEST*** | | | | | | |
| No significant responses | | | | | | |
| **3. Main effect of practice: Regression with MEP changes** | | | | | | |
| ***TRAINING \| RETEST \| TRAINING – RETEST*** | | | | | | |
| No significant responses | | | | | | |
| ***4.* Modulation in activity by speed: Regression with MEP changes** | | | | | | |
| ***TRAINING*** | | | | | | |
| Hippocampus | -20 | -16 | -24 | 21 | 3.05 | 0.027* |
| ***RETEST*** | | | | | | |
| Putamen | 26 | -8 | 8 | 3352 | 4.17 | 0.001* |
| | -28 | -14 | 2 | | 2.99 | 0.036 |
| Caudate | 14 | 12 | 4 | | 3.24 | 0.018* |
| ***TRAINING – RETEST*** | | | | | | |
| Caudate | 12 | 10 | 4 | 5355 | 3.79 | 0.003* |
| Putamen | 24 | -8 | 6 | | 4.05 | 0.001* |
| | -28 | -14 | 2 | | 3.38 | 0.012* |
| Hippocampus | 24 | -22 | -16 | | 3.17 | 0.022 |

*Asterisk (*) indicates significance at p<0.05 after Holm-Bonferroni correction for multiple comparison. Statistics were extracted from regions of interest without applying anatomical masks.*

**Supplemental Table S17.** Functional imaging results of the regression analyses between connectivity maps and overnight macro-offline gains in accuracy as well as changes in MEPs – main effect of group (F test)

| Area | x | y | z | k | Z | $p_{FWEsvc}$ |
|---|---|---|---|---|---|---|
| **1. Regressions with macro-offline gains in accuracy** | | | | | | |
| **1.1 TRAINING** | | | | | | |
| ***Hippocampus [-28 -10 -26] connectivity*** | | | | | | |
| No significant responses | | | | | | |
| ***Caudate [-8 16 4] connectivity*** | | | | | | |
| No significant responses | | | | | | |
| ***DLPFC target [-30 22 48] connectivity*** | | | | | | |
| No significant responses | | | | | | |
| ***Modulation in DLPFC target [-30 22 48] connectivity by speed*** | | | | | | |
| No significant responses | | | | | | |
| **1.2 TRAINING – RETEST** | | | | | | |
| ***Caudate [-6 12 6] connectivity*** | | | | | | |
| No significant responses | | | | | | |
| **2. Regressions with MEPs changes** | | | | | | |
| **2.1 TRAINING** | | | | | | |
| ***Hippocampus [-28 -10 -26] connectivity*** | | | | | | |
| No significant responses | | | | | | |
| ***Caudate [-8 16 4] connectivity*** | | | | | | |
| Hippocampus | 28 | -28 | -6 | 117 | 3.26 | 0.015* |
| | -24 | -34 | -4 | 31 | 2.97 | 0.034* |
| ***DLPFC target [-30 22 48] connectivity*** | | | | | | |
| No significant responses | | | | | | |
| ***Modulation in DLPFC target [-30 22 48] connectivity by speed*** | | | | | | |
| No significant responses | | | | | | |
| **2.2 TRAINING – RETEST** | | | | | | |
| ***Caudate [-6 12 6] connectivity*** | | | | | | |
| Hippocampus | -16 | -6 | -22 | 21 | 2.92 | 0.04* |

*Asterisk (*) indicates significance at p<0.05 after Holm-Bonferroni correction for multiple comparison. Statistics were extracted from regions of interest without applying anatomical masks.*

### 4 Supplemental Figures

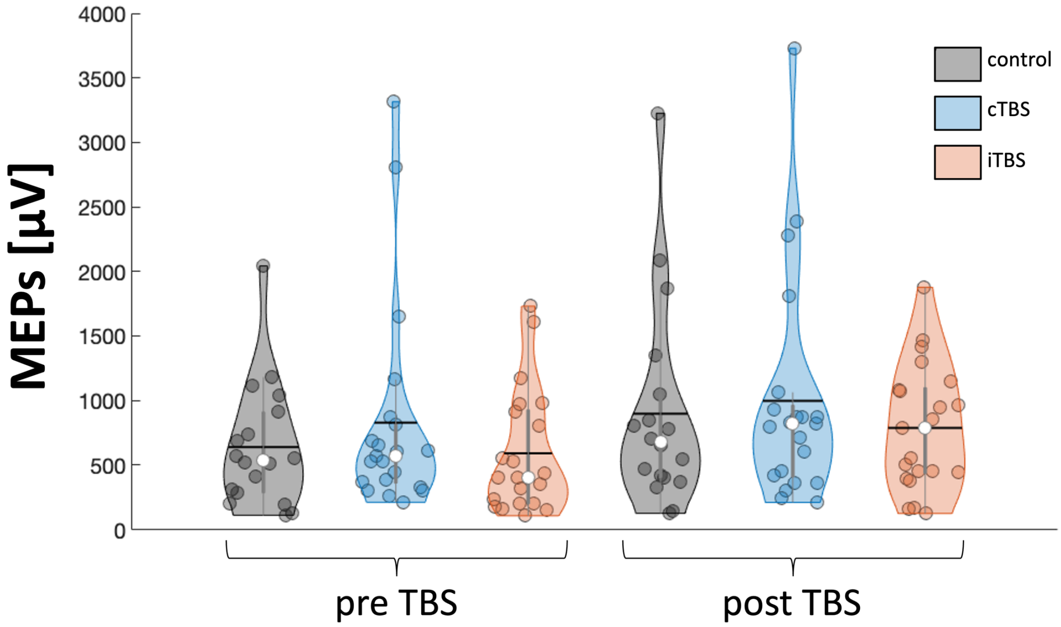


Supplemental Figure S1. MEP values. Raw MEP values increased from pre- to post-TBS similarly in all three groups. Colored circles represent individual data, jittered in arbitrary distances on the x-axis within the respective violin plot to increase perceptibility. Black horizontal lines represent means and white circles represent medians. The shape of the violin plots depicts the distribution of the data and grey vertical lines represent quartiles. MEPs: motor evoked potentials, TBS: theta-burst stimulation, c: continuous, i: intermittent.

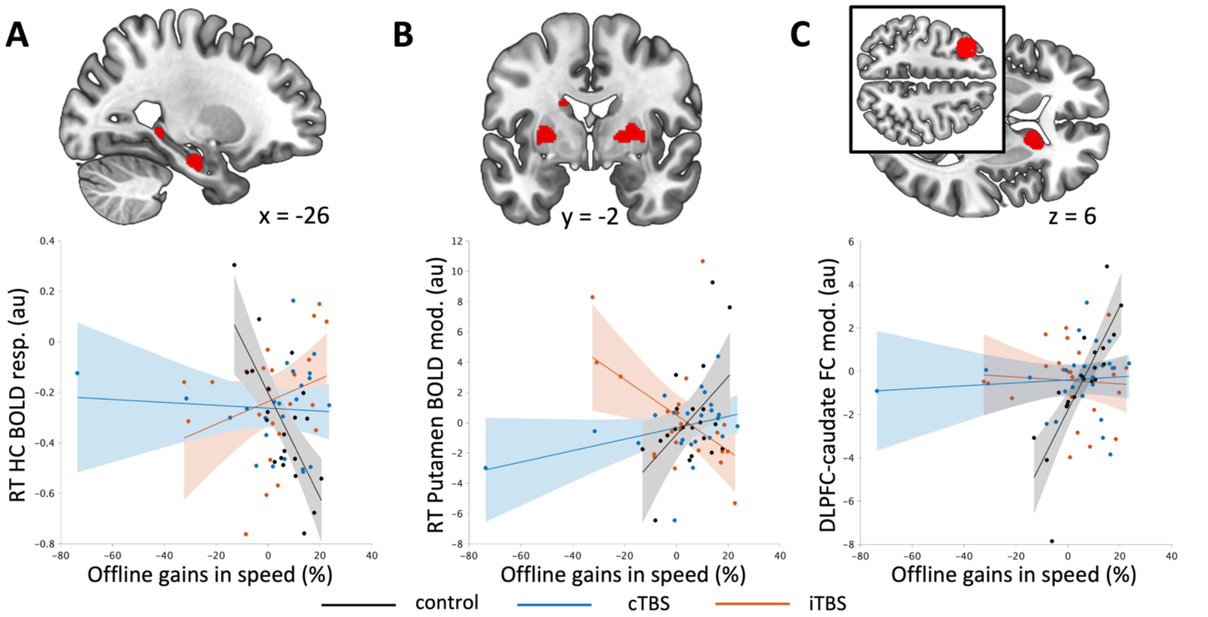


<u>Supplemental Figure S2.</u> Main effect of group on the relationship between task-related responses and macro-offline gains in performance speed. (A) Between-group differences in the relationship between task-related hippocampal (HC) activity (anterior and posterior parts) during retest and macro-offline gains in speed. Higher hippocampal activity during inter-practice rest intervals (i.e., more negative beta values) during retest was related to greater macro-offline gains in performance in the control group compared to the active stimulation groups. (B) Between-group differences in the relationship between macro-offline gains in performance speed and dynamical activity in the striatum during retest. Macro-offline gains in performance were associated to a progressive decrease in activity as a function of practice during retest in the control group (higher beta values represent more block-to-block decrease in activity) while it was related to a practice-related increase in the iTBS group. Correlation differed significantly between the control group and the two active stimulation groups, as well as between the two active stimulation groups (Supplemental Table S13). (C) Between-group differences in the relationship between offline gains in performance speed and dynamical functional connectivity (FC) between the DLPFC (seed region depicted in black box) and the caudate nucleus during training. Offline gains in performance were associated to a progressive decrease in DLPFC-caudate connectivity as a function of practice during training in the control group (higher beta values represent more block-to-block decrease in connectivity) compared to the active stimulation groups. Correlation differed significantly between the control group and the two active stimulation groups (Supplemental Table S15). Regression maps are displayed in the ROIs on a T1-weighted template image at a threshold of $p<.005$, uncorrected. Circles represent individual data, solid lines represent linear regression fits, shaded areas depict 95% prediction intervals of the linear function. Resp.: response, au: arbitrary unit, mod: modulation contrast, DLPFC: dorsolateral prefrontal cortex, cTBS: continuous theta-burst stimulation, iTBS: intermittent theta-burst stimulation.
